# *In situ* structure of rotavirus VP1 RNA-dependent RNA polymerase

**DOI:** 10.1101/605063

**Authors:** Simon Jenni, Eric N. Salgado, Tobias Herrmann, Zongli Li, Timothy Grant, Nikolaus Grigorieff, Stefano Trapani, Leandro F. Estrozi, Stephen C. Harrison

**Affiliations:** Department of Biological Chemistry and Molecular Pharmacology, Harvard Medical School, 250 Longwood Avenue, Boston, MA 02115, USA; Laboratory of Molecular Medicine, Boston Children’s Hospital, Boston, MA 02115, USA; Graduate Program in Virology, Harvard Medical School, Boston, MA 02115, USA; Janelia Research Campus, Howard Hughes Medical Institute, Ashburn, VA 20147, USA; Univ. Montpellier, Centre de Biochimie Structurale (CBS), INSERM, CNRS, 34090 Montpellier, France; Univ. Grenoble Alpes, CNRS, CEA, Institut de Biologie Structurale (IBS), 38000 Grenoble, France; Howard Hughes Medical Institute, Harvard Medical School, Boston, MA 02115, USA

**Keywords:** non-enveloped virus, viral RNA-dependent RNA polymerase, RNA transcription, local reconstruction, electron cryomicroscopy (cryo-EM)

## Abstract

Rotaviruses, like other non-enveloped, double-strand RNA (dsRNA) viruses, package an RNA-dependent RNA polymerase (RdRp) with each duplex of their segmented genomes. Rotavirus cell entry results in loss of an outer protein layer and delivery into the cytosol of an intact, inner capsid particle (the “double-layer particle” or DLP). The RdRp, designated VP1, is active inside the DLP; each VP1 achieves many rounds of mRNA transcription from its associated genome segment. Previous work has shown that one VP1 molecule lies close to each fivefold axis of the icosahedrally symmetric DLP, just beneath the inner surface of its protein shell, embedded in tightly packed RNA. We have determined a high-resolution structure for the rotavirus VP1 RdRp *in situ*, by local reconstruction of density around individual fivefold positions. We have analyzed intact virions (“triple-layer particles” or TLPs), non-transcribing DLPs and transcribing DLPs. Outer layer dissociation enables the DLP to synthesize RNA, *in vitro* as well as *in vivo*, but appears not to induce any detectable structural change in the RdRp. Addition of NTPs, Mg^2+^, and S-adenosyl methionine, which allows active transcription, results in conformational rearrangements, in both VP1 and the DLP capsid shell protein, that allow a transcript to exit the polymerase and the particle. The position of VP1 (among the five symmetrically related alternatives) at one vertex does not correlate with its position at other vertices. This stochastic distribution of site occupancies limits long-range order in the 11-segment, dsRNA genome.

## Introduction

Rotaviruses, like other dsRNA viruses, encapsidate RNA-dependent, RNA polymerases (RdRps) to transcribe their segmented genomes. In the infectious virion, known as a “triple-layer particle” (TLP), three concentric, icosahedrally symmetric protein shells surround the packaged RNA (Fig. 1a) [1,2]. The role of the outer layer, composed of viral proteins VP4 and VP7, is to introduce the virion into a cell and to deliver into the cytosol the double-layer particle (DLP) that it surrounds. The proteins of the DLP include VP2 and VP6, as well as the RdRp (VP1) and a capping enzyme (VP3). Loss of the outer layer during entry (or from its removal *in vitro*) derepresses the transcriptional activity of VP1 and the capping activity of VP3 within the DLP. The structure of VP2 and its organization in a 120-subunit inner shell are characteristics of nearly all dsRNA viruses [3]. Because the viral protein numbering varies, depending on the class to which the virus belongs, we refer here to VP2 and its homologues (as have other authors) as the capsid-shell protein (CSP). The DLP interior contains one copy of the VP1 RdRp for each of the 11 dsRNA genomic segments and a comparable, but still undetermined, number of VP3 subunits.

**Fig. 1.**
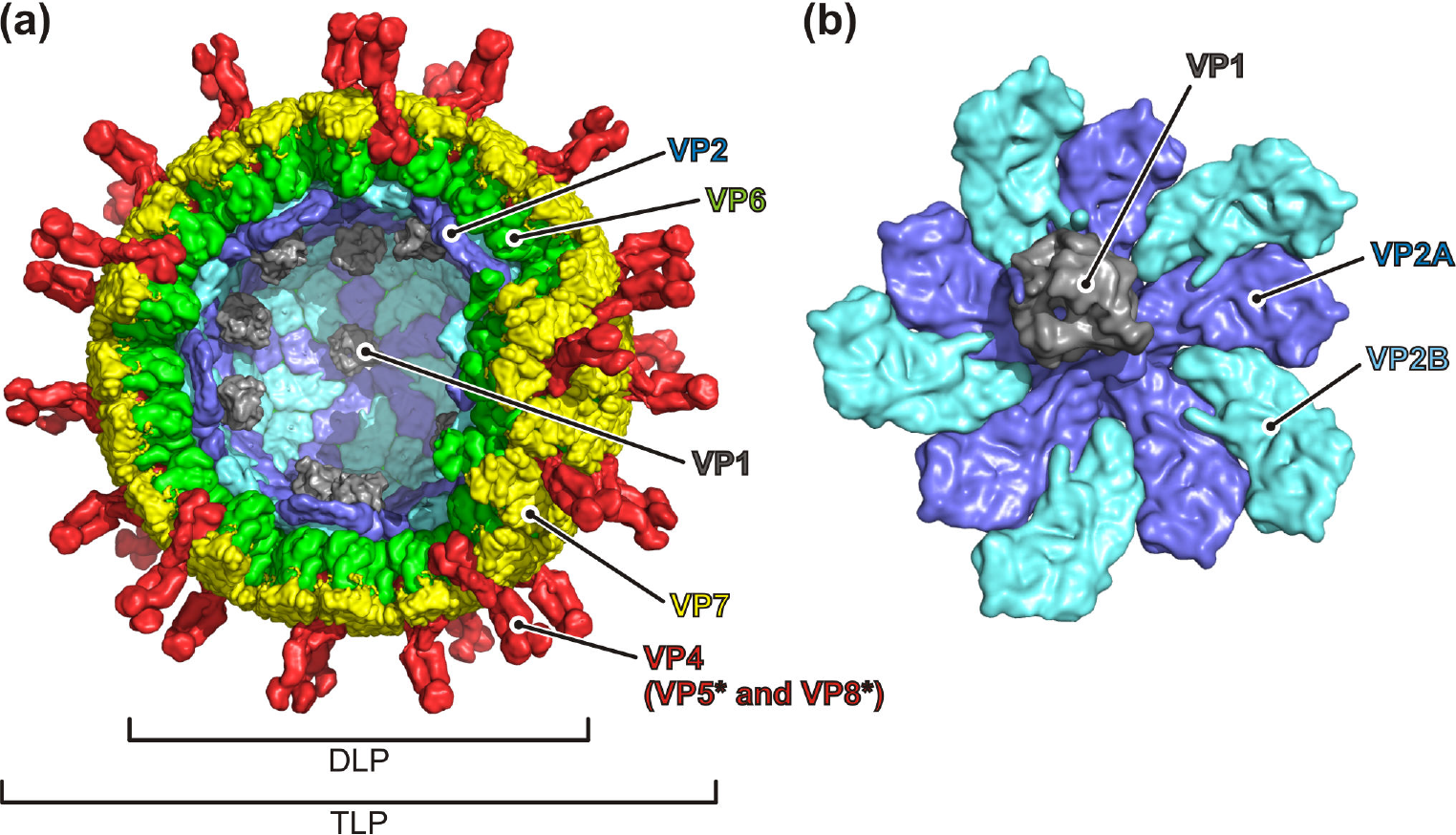
Overview of the rotavirus architecture. (a) “Triple-layer particle” (TLP) in surface representation with part of the capsid removed to allow inside view (VP1, gray; VP2A, blue; VP2B cyan; VP6, green; VP7, yellow; VP5* and VP8*, red). (b) Close-up view of a fivefold vertex surrounded by VP2 subunits (blue and cyan) and one bound VP1 polymerase (gray).

RdRps of dsRNA viruses transcribe mRNA conservatively from dsRNA genomic segments; they also copy the minus-sense strand of those segments *in situ*, from the plus-sense strands packaged during assembly of the capsid shell. The resulting, transcriptionally active DLPs can also be the source of further plus-sense strands, until the production of viral proteins switches to packaging them into TLPs. Crystal structures of reovirus and rotavirus RdRps show that the active site lies within a cage-like surround, with access channels for template and nucleoside triphosphate (NTP) substrates and exit channels for transcript and template (or dsRNA product in the replication step) [4–6]. To achieve this cage-like organization, an N-terminal domain and a C-terminal domain augment a familiar fingers-palm-thumb polymerase domain. The largely α-helical, C-terminal domain surrounds the channel for template or nascent genome exit -- hence, its designation as the “bracelet domain” (surrounding the wrist of the fingers, palm and thumb).

In earlier work, we showed that in DLPs, VP1 associates with the inner surface of the VP2 capsid shell, at positions overlapping each of the fivefold axes [7]. Even in the averaged density of an icosahedrally symmetric reconstruction from cryo-EM images, part of the protein projected far enough from the fivefold axis that we could place its crystallographically determined structure by fitting the non-overlapping features of the map (Fig. 1b) [7]. Since then, asymmetric reconstructions of cytoplasmic polyhedrosis virus (CPV) and of an aquareovirus (ARV) have yielded full, *in situ* structures of their RdRp enzymes and of a closely associated NTPase [8,9]. Like mammalian orthoreoviruses, CPV and ARV have external, turret-like structures, composed of the capping-enzyme protein, projecting along each of their fivefold axes. Transcribed RNA emerges through these turrets, which have guanylyl transferase and methylase activities. A likely function of the internal NTPase is to ensure that the nascent transcript lacks a 5’-γ-phosphate, so that transfer of GMP by the guanylyl transferase yields the standard 3’-GpppG-3’ cap structure; the turrets themselves do not appear to have such an activity. The orientation of the rotavirus VP1 RdRp with respect to the capsid shell suggests that the transcript could pass directly from the transcript exit channel into an opening at the fivefold axis. The same is true for CPV and ARV, allowing direct passage of the nascent transcript into the capping chamber of the turret. Because rotaviruses extrude fully capped transcripts, with all relevant enzymatic activities incorporated into VP3, their internal organization must allow this order of events.

We describe here high-resolution structures for the rhesus rotavirus (RRV) VP1 RdRp *in situ*, achieved by local reconstruction of density in relevant volumes around individual fivefold positions on the interior of the capsid shell. We have analyzed TLPs, non-transcribing DLPs and transcribing DLPs. The transition from TLP to DLP does not by itself appear to induce any detectable structural change in the RdRp, but addition of NTPs, Mg^2+^, and S-adenosyl methionine (SAM) leads to synthesis of RNA and to local conformational changes in the bracelet domain and in the CSP that allow a transcript to exit both the polymerase and the particle. We can visualize template and transcript in the active-site cavity of the RdRp. The position (among the five symmetrically related possibilities) of an RdRp at any particular vertex does not correlate with the orientation of RdRps at other vertices. This stochastic distribution of site occupancies appears to limit long-range order in the dsRNA genome. We can nonetheless detect density corresponding to genomic RNA and determine some of its general features.

## Results

### Cryo-EM reconstructions of VP1 RdRp within rotavirus particles

Asymmetric reconstruction gave no evidence of a unique or symmetrical arrangement of VP1 with respect to the icosahedral symmetry, and we show below that the distribution of VP1 molecules has none of the symmetries that would be a subgroup of the icosahedral point group (e.g., the pseudo-D3 symmetric distribution found in cytoplasmic polyhedrosis virus and aquareovirus [8–10]). We conclude that VP1 binds stochastically at one of the five possible positions at each of the fivefold vertices. The number of possible particle configurations is therefore around 10^8^. Classification and averaging of entire virus particle images with identical VP1 configurations is obviously not possible, and we analyzed vertices individually instead. For local reconstruction of the VP1 polymerase, we used current methods in cryo-EM data processing, including signal subtraction [11], sub-particle extraction [12], and classification [13], as described in detail in Materials and Methods (Fig. 2). After initial reference-based alignment of entire viral particles with icosahedral symmetry imposed, we extracted subparticles from the 2D images at potential VP1 sites and determined VP1 positions by classification with the help of class-specific masks. We obtained local reconstructions for rotavirus particles before (TLP and DLP) and after incubation with NTPs and SAM (TLP_RNA and DLP_RNA) (Table S1). The estimated resolution for the local reconstructions ranges from 5.2 to 3.3 Å, with the highest resolution for the DLP map (Table S1, Fig. S1), in which amino-acid side chains are resolved (Fig. S2). The resolution of VP1 density was only slightly lower than that of the viral shell (Fig. S1). Therefore, VP1 binds tightly and specifically to the inward-facing surface of VP2. We built molecular models, consisting of one VP1 molecule and ten VP2 molecules (VP2A 1–5 and VP2B 1–5) into the cryo-EM densities (Table S2). Statistics for the refined structures are in Table S1. Fig. S1 shows their correlation with the cryo-EM density.

**Fig. 2.**
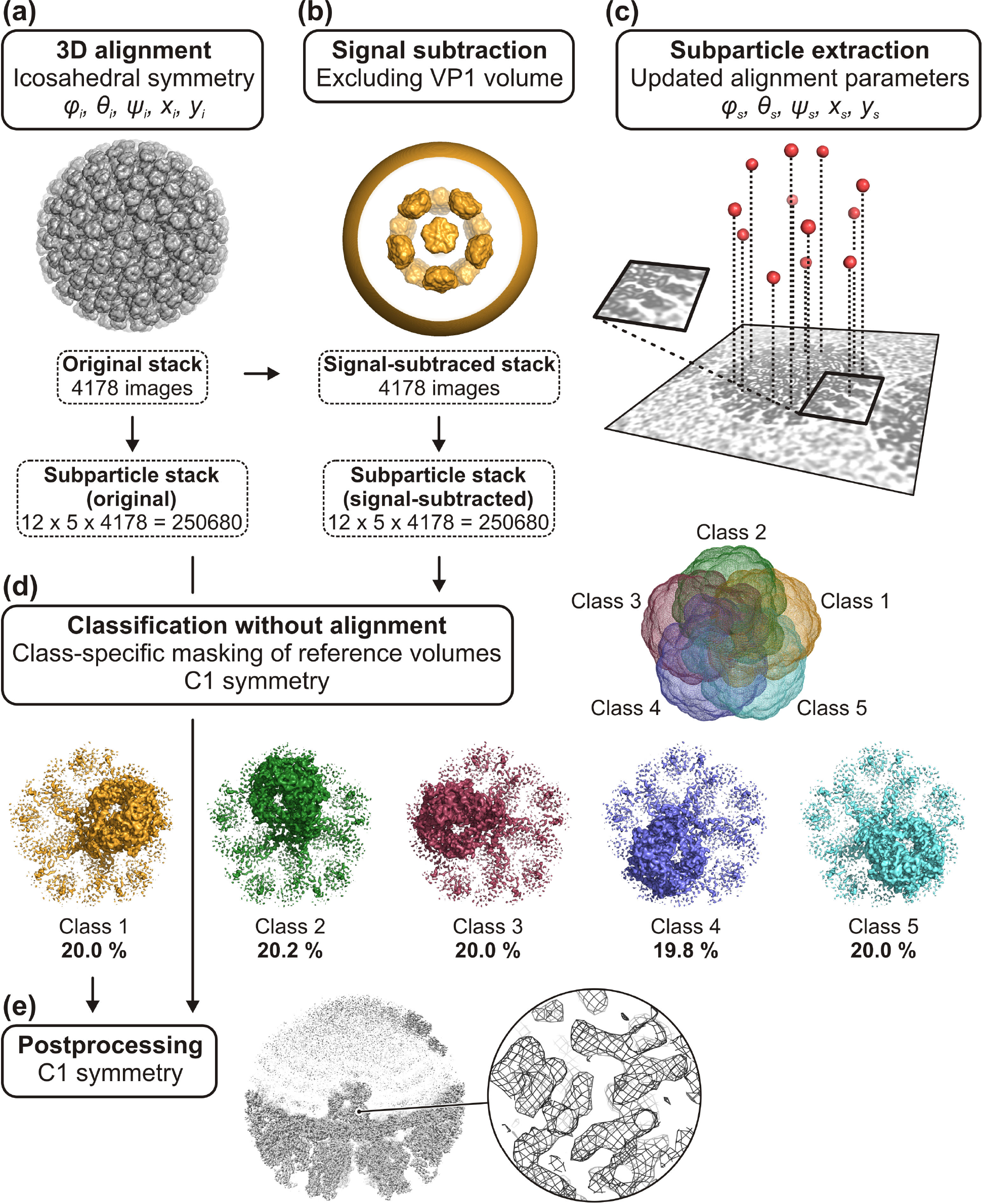
Local reconstruction of VP1 polymerase by cryo-EM (see also Materials and Methods). (a) Initial alignment of virus particles with icosahedral symmetry imposed. (b) Signal subtraction in the 2D particle images using corresponding reference projections after applying a mask that excludes density within regions potentially occupied by VP1. (c) Subparticle extraction based on the virion particle alignment parameters and icosahedral symmetry (60 subparticles per virus). Updated alignment parameters are calculated for the subparticles to reproject them on the same asymmetry for reconstruction of the subvolume. (d) Classification with class-specific masks. (e) The final map is calculated with a subparticle stack extracted from the non-signal-subtracted (original) images.

As a further check on the absence of correlations among positions of different VP1s within a particle, we recorded, for each particle, which of the five “choices” of positions was occupied by VP1 at each of the eleven vertices related to an arbitrary initial vertex by a defined permutation of icosahedaral transformations. Table S3 shows that the results are consistent with a fully random distribution of the remaining eleven with respect to the first.

### Molecular structure of VP2-bound RdRp

Rotavirus RdRp lies just inside the VP2 shell with an offset from the fivefold symmetry axis. It makes contacts with several of the surrounding VP2 molecules (Fig. 3a). From crystal structures of VP1 alone and its complexes with substrates [4], we assign VP1 residues to domains as follows: N-terminal domain, 1–332; fingers, 333–488 and 524–595; palm, 489–523 and 596– 685; thumb, 686–778; C-terminal, bracelet domain, 779–1088 (Fig. 3a). The domains fold together into a cage-like structure with substrate entry and exit tunnels leading to and from the catalytic site located in the central cavity. In the capsid-attached RdRp, the transcript-exit site faces the VP2 shell; the template-entry channel is on the side facing inward; the nucleotide-entry and template-exit channels face laterally (Fig. 3b). Our structures generally confirm our previously reported rigid-body fit of a VP1 crystal structure into icosahedrally averaged maps of rotavirus particles [7].

**Fig. 3.**
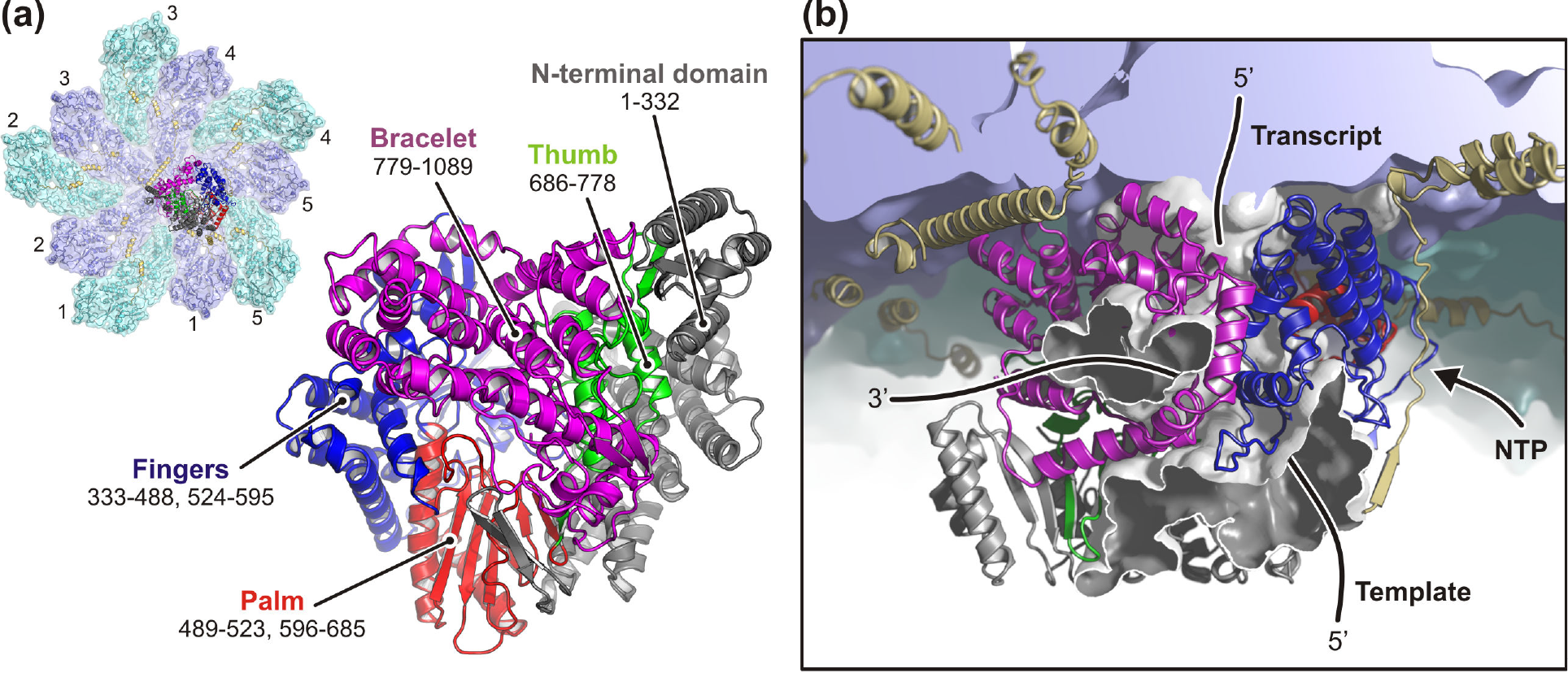
Structure of the rotavirus VP1 RNA-dependent RNA polymerase (RdRp) bound to VP2 capsid proteins (refined model from DLP dataset). (a) Inset, overview of RdRp bound to a five-fold vertex. The VP2A (blue) and VP2B (cyan) protomers are numbered 1–5. The close-up view shows the RdRp structure in conventional orientation. The N-terminal domain is colored gray; the palm subdomain is in red; the fingers, blue; the thumb, green; the bracelet, magenta. Residue numbers are the domain boundaries. (b) Side view of RdRp and the VP2 shell. The RdRp tunnel system is shown and template, transcript and nucleotides entry and exits sites, respectively, are schematically indicated.

The absence of any RNA density in the RdRp active site in the TLP or DLP reconstructions suggest that we observe the polymerase in a pre-initiation or initiation-competent state. RdRps of rotavirus replication intermediates initially bind a conserved sequence at the 3’ ends of the +RNAs (3’CS+) [14], and VP1 crystals soaked with 3’CS+ oligonucleotides show bases in the active site and the consensus sequence UGUG bound with sequence-specific contacts in the template-entry channel [4]. Packaging into the VP2 shell initiates synthesis of the complementary, minus-sense RNA strand, generating the double stranded genome. In our structures, which contain fully double-stranded RNA, we do not see specific RNA contacts in the template channel, but we can detect density just outside the polymerase, near the template entry site (Fig. S3). We suggest that it comes from weakly bound 3’ ends of -RNA strands, the template for mRNA (+RNA) production.

Superposing the fingers, palm and thumb core domain residues from the VP1 crystal structure (RdRp in isolation) [4] and from our DLP structure (VP2-bound RdRp) shows conformational changes in both N- and C-terminal domains after binding to the capsid (Fig. S4). Most shifts in the N-terminal domain are in regions with relatively high temperate factors and may not be relevant for regulating polymerase activity (Fig. S5). The one difference in the N-terminal domain that appears functionally relevant is flipping inward of a hairpin, residues 261–271, which would, in its crystal-structure conformation, clash with VP2A molecule 1. This flip in turn requires the short α helix (residues 499–508) at one end of the proposed “priming loop” [5] to unfold into a more extended conformation (Fig. 4a). In the C-terminal bracelet domain, contact with VP2 molecule 5 shifts an α-helical subdomain (residues 918–1006) up to about 8 Å from its position in the apoenzyme crystal structure (Fig. 4b). In the transcribing DLP we see a shift in the same subdomain, back to a position close to the one it has in the crystal structure.

**Fig. 4.**
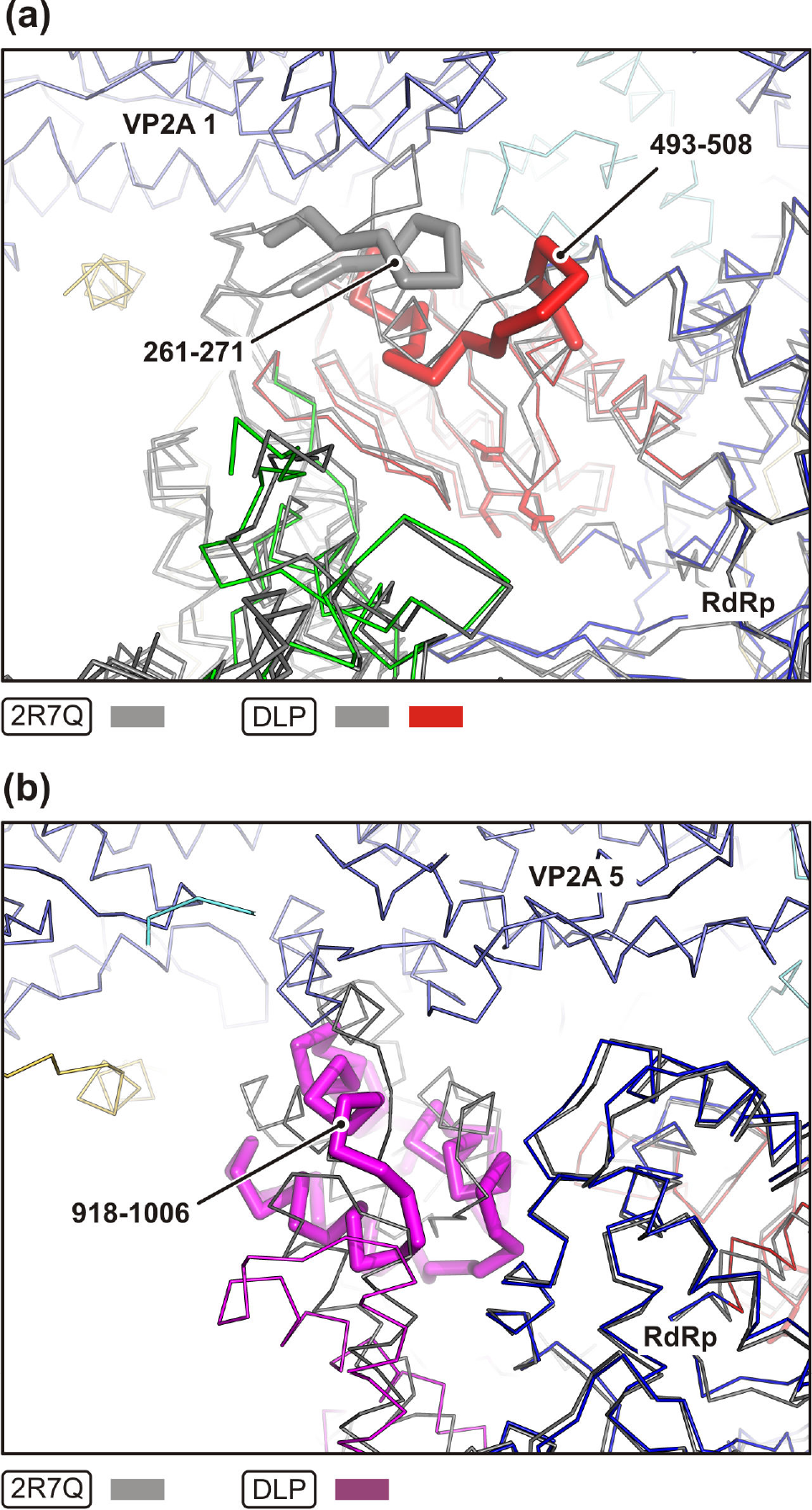
Conformational changes between the VP1 crystal structure (RdRp in isolation, colored in gray, PDB-2R7Q) and our DLP structure (VP2-bound RdRp, colored as in Fig. 3). The two structures were superimposed using corresponding residues of the fingers, palm and thumb domains only. (a) Close-up view showing the hairpin, residues 261–271, of the RdRp N-terminal domain, and the proposed “priming loop”, residues 493–508. RdRp active site residues are shown as sticks. For clarity, the bracelet domain is omitted. (b) Close-up view of the bracelet alpha-helical subdomain, residues 918–1006.

### Multiple interactions between VP1 and its VP2 neighbors

Rotavirus VP1 RdRp interacts with several VP2 capsid subunits. The enzyme binds the VP2 shell with complementary surface interactions as well as with the help of three long N-terminal VP2 extensions, which form tentacle-like interactions with VP1 (Fig. 5a). Deletion of these N-terminal extensions eliminates VP1 incorporation into recombinant virus-like particles but does not prevent capsid-shell assembly [15]. Except for the tentacle interactions, the largest interface is with VP2A molecule 1, burying a solvent accessible surface of 1622 Å^2^ (DLP structure). A long β hairpin from the VP1 N-terminal domain (residues 258–275), loops of the palm domain, and part of the bracelet domain together accommodate the tip of loop 361–372 on VP2A molecule 1. Contacts between VP2B molecule 5 and the palm and finger of VP1 and between VP2A molecule 5 and the finger and bracelet domains have buried surfaces of 1065 Å^2^ and 1047 Å^2^, respectively. At the latter contact, a helix of the bracelet subdomain packs against the tip of VP2A molecule 5 (residues 345–374). Modulation of the interaction between these two α helices appears to be important for transcription and RNA extrusion from the particle, as we discus in more detail below; the tip of VP2A molecule 5 appears to be a “gate” for RNA exit. VP2A molecules 2, 3 and 4 form additional minor contacts with VP1. The various contacts summarized here lock the RdRp firmly into place, as reflected by the relatively low temperature factors in the core of the enzyme (Fig. S5).

**Fig. 5.**
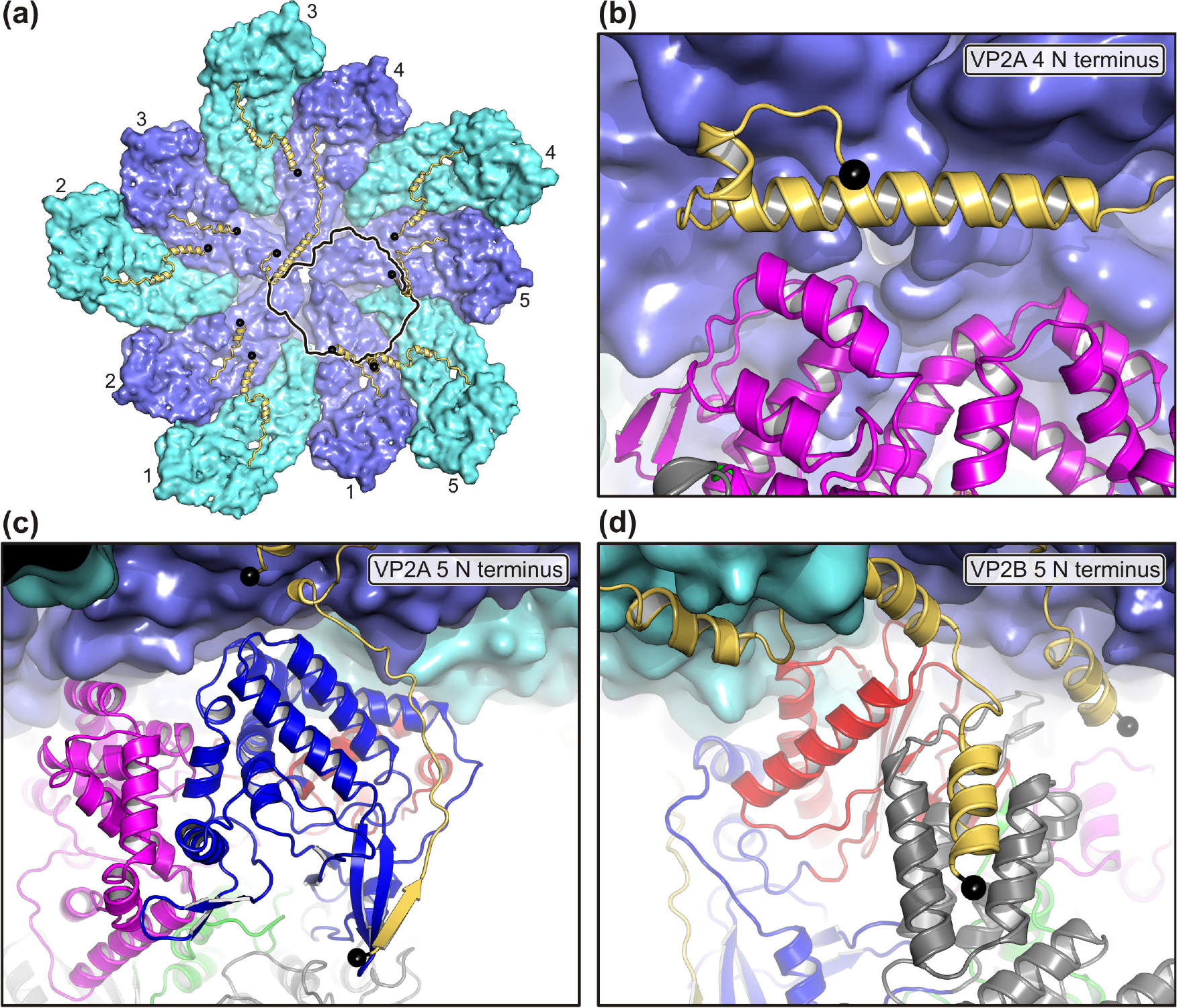
Interactions between VP1 and VP2 (refined model from DLP dataset). (a) The VP2A (blue) and VP2B (cyan) protomers are numbered 1–5 and shown in surface representation, except their N termini, which are colored yellow and are shown in ribbon representation. The first residues visible in the cryo-EM map are indicated by black spheres. The footprint of RdRp is shown as black line. (b) The VP2A 4 N terminus (tentacle 1) folds into a long α helix that interacts with the RdRp bracelet domain. (c) The VP2A 5 N terminus (tentacle 2) forms a long extension that interacts at its tip with a β hairpin of the RdRp finger domain. (d) The VP2B 5 N terminus (tentacle 3) folds into a short α helix that interacts with the RdRp N-terminal domain.

Tentacle 1 is the N-terminal arm of VP2A molecule 4; it folds into a long α helix that lies in a groove between the capsid shell and the VP1 bracelet domain (Fig. 5b). Tentacle 2, the N-terminal arm of VP2A molecule 5, wraps in an extended conformation around the RdRp and forms a short strand with a β hairpin of the fingers (Fig. 5c). Tentacle 3, the N-terminal arm of VP2B molecule 5, packs with a short α helix against the RdRp N-terminal domain (Fig. 5d). The conformational variability of the VP2 N-terminal arm (Fig. S6) facilitates the multiple interactions with a single, asymmetrically placed VP1 molecule. Because one RdRp binds all five VP2A and one VP2B molecule of a 5-fold capsid hub with either interface- or tentacle-mediated contacts, VP1 might enhance stability of VP2 decamers during virus maturation and plus-strand packaging.

### Structural differences between TLP and DLP

Transcriptionally competent DLPs are released into the cytoplasm after the VP7 layer dissociates from the inner capsid [16]. Structural differences between VP7-coated TLPs and uncoated DLPs might therefore in principle explain how loss of the outer layer activates the RdRp. Comparison of icosahedrally averaged TLP and DLP structures showed a distinct deformation of the VP2-VP6 fivefold hub, associated with a previously described shift in orientation of the VP6 trimers around the fivefold axis and a concomitant shift of the underlying, fivefold proximal tips of VP2A [17,18]. We see the same difference in the locally reconstructed maps: in the DLP structure, the tips of VP2A tilt outwards, displacing the loop closest to the fivefold axis by about 6 Å from its position in the TLP structure (Fig. S7). The diameter of narrowest section in the transcript exit pore, formed by the tips of the VP2A molecules (residues 346–373) remains the same, however, ruling out the suggestion that straightforward pore opening might regulate activation of transcription in DLPs [18]. Except for outward displacement of the VP2A tips and an accompanying displacement of the RdRp, molecule-by-molecule superpositions in our current structures show that neither VP2 nor VP1 undergoes any major conformational change in the transition from TLP to DLP (Fig. S8).

### Molecular structure of a transcribing RdRp

To visualize structures of transcribing rotavirus particles, we incubated TLPs and DLPs with nucleotides and S-adenosylmethionine (SAM) before freezing grids for cryo-EM analysis. TLP images occasionally showed a transcript strand emerging from particles, but no accumulation of released RNA in the background (TLP_RNA in Fig. S9a). The RdRp reconstruction calculated from this TLP_RNA sample showed an empty active site, with no density for template or transcript, and no substantial conformational differences in VP1 from its structure in the DLP. Thus, most polymerases in these TLPs did not transcribe when nucleotides were added, consistent with many published studies.

In the images of DLPs incubated with nucleotides and SAM, almost all particles inspected showed transcript strands emerging from the capsids, as previously reported [19], and single stranded RNA molecules accumulated in the background (DLP_RNA in Fig. S9b). The RdRp reconstruction calculated from this DLP_RNA sample showed template and transcript density in the active site. Because we did not attempt to stall RdRp at a specific template position before freezing grids, and because the RNA sequences of the various genome segments are different, the reconstruction is an average of polymerases captured at different transcriptional positions. We therefore used generic RNA sequence to fit the reconstruction. We built into the observed density a ten-nucleotide long template-transcript double helix in the cavity of the RdRp and four additional nucleotides (positions n+1 to n+4) in the template-entry channel. The density for template nucleotides in the entry channel is of relatively low resolution, consistent with the ensemble of sequences present there. Both template and transcript densities become well defined where the minus-strand template bends to form base pairs with the incoming (position n) and priming (position n-1) nucleotides. After nine base pairs, the template-transcript double helix splits into single strands. Blurred density shows the template strand exiting thought the central aperture of the bracelet domain and the transcript strand leaving the RdRp between the bracelet and fingers domains.

### Structural differences between non-transcribing and transcribing DLPs

There are notable conformational changes in VP1 between initiation-competent and transcriptionally active DLPs (Fig. S10). In order to accommodate the template-transcript RNA double helix within the RdRp cage, the loop between residues 486–507 (the proposed priming loop) and the tip of the hairpin 261–271 in the N-terminal domain shift to make space for the growing duplex. As seen in the crystal structures [4], loop 397–404 of the fingers domain also retracts from the central cavity, to allow the template to move into the catalytic site. Smaller shifts occur in residue ranges 575–582, 629–632 and 818–827. Residues 918–1006, an α-helical subdomain of the bracelet, shows the most substantial displacement in the transcribing DLP structure, shifting by up to 10 Å outwards from its position in the non-transcribing DLP. This rearrangement allows passage of the transcript strand, with which that subdomain would otherwise clash (Fig. 6). The subdomain is at the position at which the template and transcript strands separate. The C-terminal plug (residues 1074–1088), which would interfere with elongation, has withdrawn from the central cavity, and the last residue for which we observe density, is 1073. Side-chain contacts between the C-terminal plug and the bracelet subdomain may restrain the latter, so that displacement of the plug as the emerging template pushes against it might then allow the subdomain to move away more freely.

**Fig. 6.**
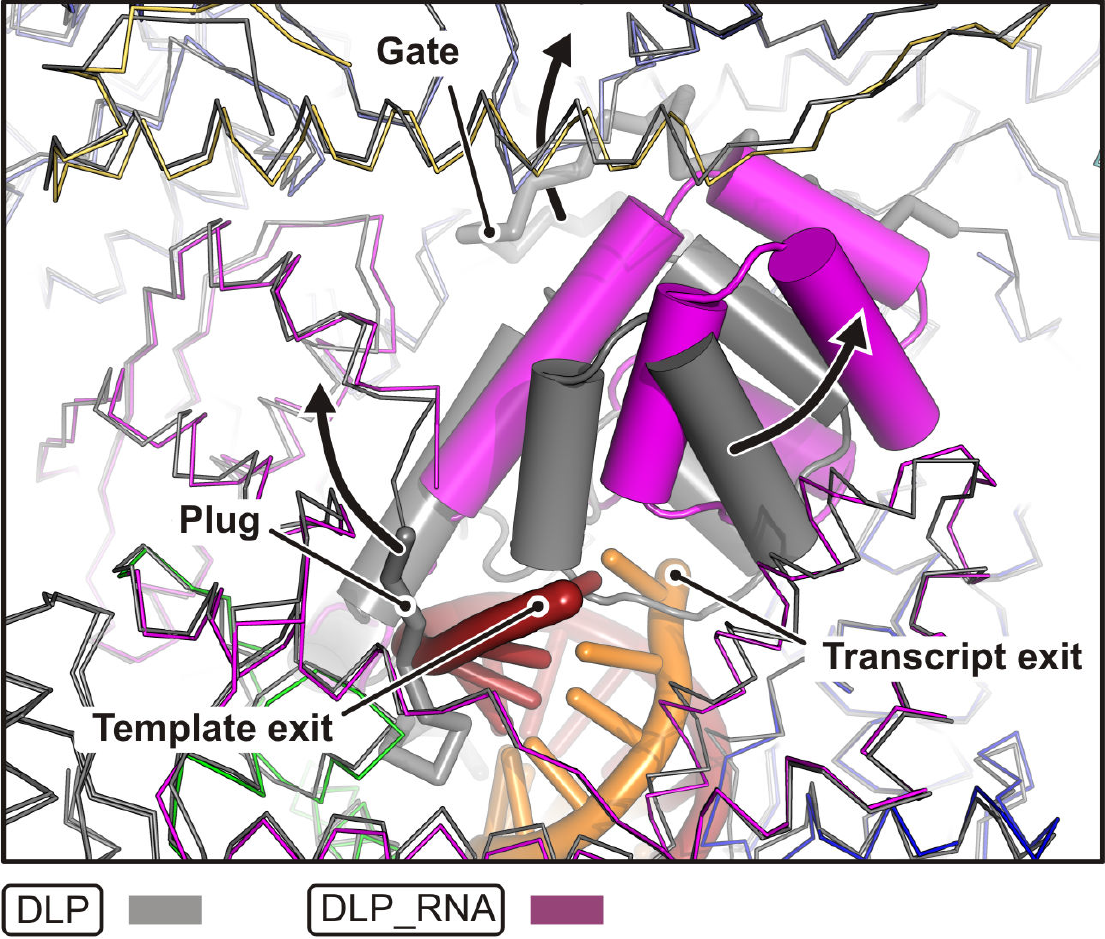
Rotavirus VP1 RdRp structure of transcriptionally active DLPs (refined models from DLP and DLP_RNA datasets). Close-up view centered on the RdRp template exit tunnel. The DLP_RNA structure (shown in color) is superposed on the DLP structure (shown in gray). The DLP_RNA RdRp and VP2 molecules are colored as in Fig. 3. The bracelet domain is in magenta; the template RNA, dark red; the transcript RNA, orange. Arrows indicate major conformational changes between DLP and DLP_RNA that occur in RdRp and VP2 upon incubation with nucleotides: (i) a large movement of part of the bracelet domain, (ii) release of the C-terminal plug form the active site, (iii) release of the VP2A molecule 5 gate that allows access of the transcript to the five-fold vertex pore.

Rearrangement of the α-helical bracelet subdomain (residues 918–1006) changes its interface with the capsid VP2A molecule 5 (Fig. 6). In the non-transcribing DLP structure, the ordered tip of VP2A molecule 5 (resides 346–373) blocks the RdRp transcript-exit site (Fig. 7a). Displacement of the bracelet subdomain in turn displaces these tip residues, which become too disordered to trace, creating a continuous tunnel between the RdRp central cavity and the pore on the capsid fivefold axis (Fig. 7b). We detected additional difference density (cryo-EM map – refined model) in the capsid pore of the DLP_RNA reconstruction. We interpret this density as emerging RNA transcripts, as we do not see it in the DLP reconstruction (Fig. 7c and d). The 346–373 loop of VP2A molecule 5 thus appears to be a gate, forced open by displacement of the bracelet subdomain in actively transcribing DLPs, to allow release of mRNA transcripts. We note that the conformation of this gate loop in VP2A molecule 5 is different from its conformation in VP2A molecules 1–4, because of interactions with VP1 (Fig. S6b).

**Fig. 7.**
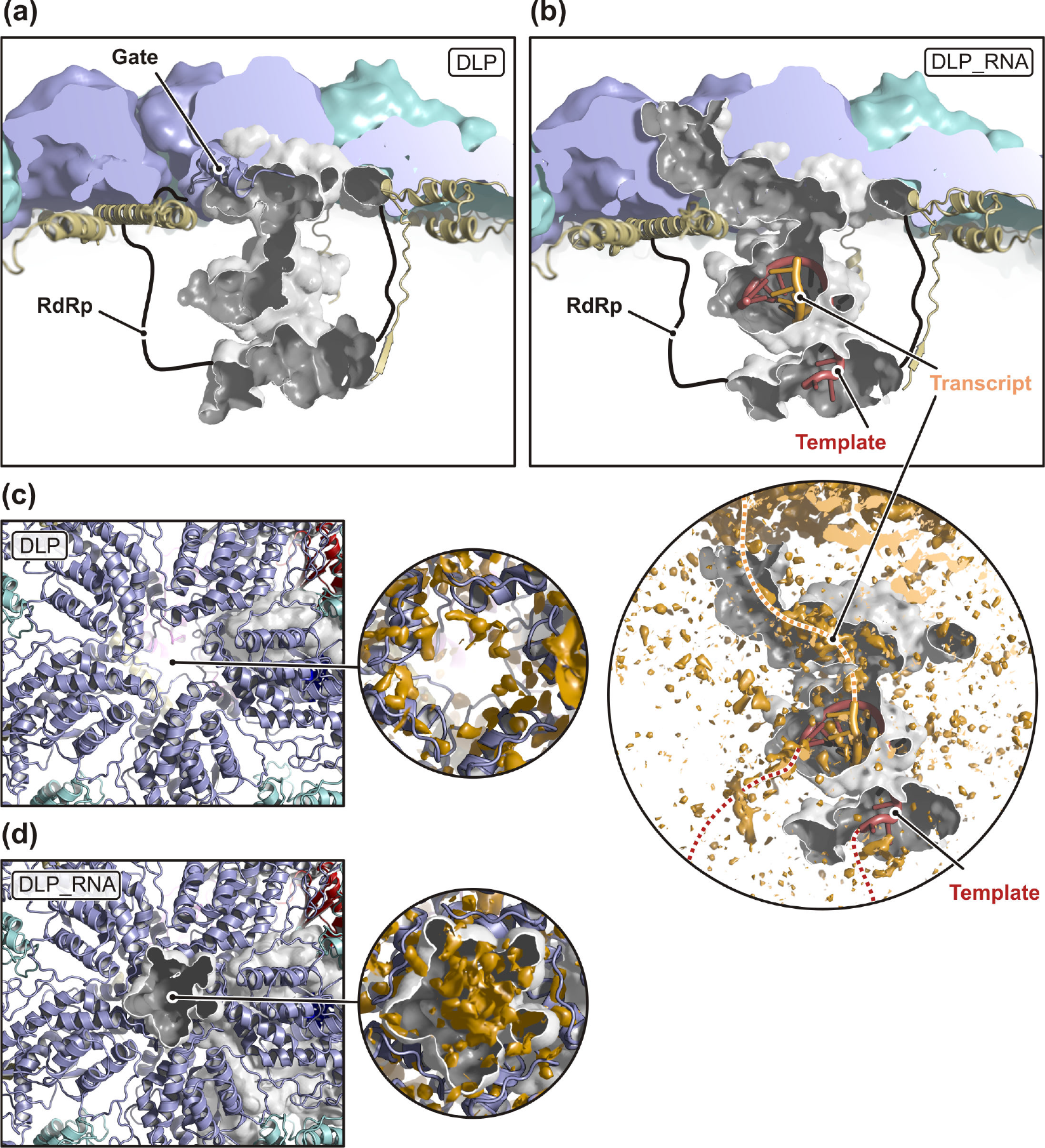
Transcript RNA exit path in transcriptionally active DLPs (refined models from DLP and DLP_RNA datasets). (a and b) Views of the RdRp internal tunnel system probed in the DLP (a) and DLP_RNA (b) structures. The VP2 shell (blue and cyan) and the tunnel (gray) are partially cut. VP2 N termini are shown as yellow ribbons. The shape of VP1 RdRp is shown schematically as outline. RNA template and transcript strands are in dark red and orange, respectively. Upon transcription, the VP2A gate becomes flexible and allows a continuous connection between the RdRp active site interior and the VP2 pore on the five-fold vertex. The magnification in (b) shows the tunnel only with difference density (cryo-EM map - model) in orange. (c and d) View of the VP2 five-fold vertex pore. The magnifications show the difference density (cryo-EM map - model) in orange for DLP (c) and DLP_RNA (d).

### dsRNA genome structure

The 11 segments of the rotavirus dsRNA genome vary in length from 0.7 to 3.3 kb. A plot of spherically averaged density shows that most of the genome packs into 8 concentric layers inside the virion with an average inter-layer spacing of 28 Å (Fig. S11). In maps aligned on the RdRps of one fivefold vertex in rotavirus DLPs or TLPs, the RNA double strands are better resolved in the outermost layer than in the others, but less so than in CPV and ARV (Fig. S12). In those turreted dsRNA viruses, the RdRp orientations correlate from vertex to vertex, enabling relatively long-range order of the dsRNA segments wound among them. In rotavirus particles, the stochastic distribution of RdRps among the five alternative positions at each vertex rules out any unique long-range order for the RNA. The substantial length variation among genome segments may also contribute to variability of even the dsRNA immediately adjacent to the RdRp. Some tendency for RNA to wrap against VP1 is evident, consistent with relatively tight overall packing within the particle, but attempts to classify individual segments or improve the resolution by focusing the alignment on RNA density did not succeed. We cannot rule out models in which the dsRNA segments spiral around the RdRp at each 5-fold vertex, but overall gentler genome bending, as found in CPV and aquareovirus, is more consistent with the persistence length in solution (Fig. S12).

There are two contacts between the bulk dsRNA density and the polymerase, likely arising from density of a bound 5’ +RNA cap structure and the corresponding 3’ -RNA strand, respectively. The first contact emerges from the putative cap-binding domain of RdRp (see below). The second contact is close to the template entry tunnel (Fig. S3), where the 3’ -RNA strand inserts to initiate transcription.

### Cap binding site

A site that binds the +RNA cap structure (m^7^GpppG) is present on the surfaces of both reovirus and rotavirus RdRps, near the opening of the template entry channel. Cap binding has been proposed to retain the 5’ end of the genomic plus-strand during transcription and to participate in recruiting plus-strand segments for packaging during virus assembly [4]. Retention of the 5’- end of the plus strand at the cap-binding site positions the 3’ end of the minus-strand near the entry of the template channel to ensure proper re-insertion for successive rounds of transcription. Without such a mechanism, the 3’ end of the minus strand would “get lost” in the particle interior after emerging from the template exit, rather than remaining poised for re-insertion into the template entry channel. The cap-binding site on reovirus λ3 was detected by soaking crystals with m^7^GpppG [5]; GTP binds a corresponding site on rotavirus VP1 between the N-terminal domain and the tip of the thumb subdomain [4]. At this position, our reconstructions show a connection between density for dsRNA and the RdRp, suggesting that the site indeed captures the capped genomic strand (Fig. S13), but the bridge of density is not well enough defined to allow modeling of molecular features.

## Discussion

### *Determining VP1 structures* in situ

We have used high-resolution cryo-EM to determine structures of the rotavirus VP1 RdRp *in situ* within the viral capsid, including its structure when actively engaged in transcription. In an initial, icosahedral reconstruction, the RdRp features are an average of five possible positions at each vertex. Tight and specific coordination of the RdRp by the capsid protein VP2 allowed us to use focused classification without alignment and hence to reduce substantially the sources of variability and noise in the final RdRp map (Fig. 2). We have found the opposite to be true of the VP5*/VP8* spikes on TLPs: flexibility (and hence a continuum of conformations) limits the resolution of spike density on TLP reconstructions [20], and our attempts to improve the maps by local alignment have been of limited success (data not shown) [12]. In the present case, imperfect signal subtraction (e.g. from the poorly ordered dsRNA genome), overlap of non-subtracted density in the subparticle images, and the reduced size of the masked structure would probably have frustrated efforts to calculate local alignments but did allow robust classification into the five discrete alternatives. The resolution we obtained for the RdRp is comparable to the resolution for VP2 (Fig. S1 and Table S1).

A very different procedure to locate the VP1 RdRp in rotavirus DLPs has been used as a test case for finding proteins or protein complexes with known high-resolution structures embedded in a dense and irregular matrix of other macromolecules (i.e., in rotavirus particles, the genomic RNA [21]). In that work, a search with a DLP “asymmetric unit” (a VP2A-VP2B pair plus 13 associated VP6 subunits) determined the positions and orientations of the sixty such units in a particle. A constrained search with the VP1 model, scanning only the immediate neighborhoods of the sixty possible location-orientation combinations, then determined the locations and orientations of nearly all eleven molecules in each of the roughly 4000 particles in that analysis. A reconstruction of the VP1 structure from the corresponding masked projected densities (roughly 15,000) had a resolution somewhat lower than ours (as assessed by inspection of the density in their posted file [Supplementary file 1, Rickgauer et al 2017] as no direct resolution estimate was given) and showed at least one of the interactions with VP2 arms that we have described here.

### RdRp distribution

The position of the RdRp at any one vertex of a rotavirus particle does not correlate with its position at other vertices (Table S3). In ARV, and probably also in orthoreoviruses, the RdRp, its partner nucleoside triphosphatase (NTPase), and the ordered regions of the eleven-segment dsRNA genome obey an approximate D3 symmetry, with one vertex devoid of the two enzymes (and with genome RNA occupying the corresponding volume). CPV, with a 10-segment dsRNA genome, has a similarly incomplete D3 symmetry, with two enzyme-less vertices. Close examination of the particle structures shows that different long-range contacts and therefore slightly different assembly pathways can account for the presence or absence of correlation among the RdRp locations in reoviruses and rotaviruses, respectively.

The likely assembly unit of all dsRNA viral capsids is a decamer of CSP subunits (in rotavirus terminology, 5 VP2A subunits and 5 VP2B subunits) together with the associated enzyme or enzyme complex. The most reasonable way to account for selection of one each of the 10-12 RNA segments (depending on the viral type in question) is to postulate that segment-segment recognition, through interactions not yet characterized, leads to a clustering of the relevant set of plus-strand segments and that the RdRp recognizes the conserved 3’ terminal +RNA sequence (the 3’CS+), thereby associating an assembly unit with each of the RNA segments. The template entrance channel of rotavirus VP1 does indeed show specific binding with a conserved tetranucleotide spaced three residues from the 3’-terminus of the plus-sense RNA strand. When bound in this way, the template “overshoots” the catalytic site by one nucleotide [4]; one interpretation of this register is that the assembly mode of RdRp-3’CS+ interaction inhibits initiation of transcription and that activation requires a one-nucleotide reverse shift of the template strand.

Differences in the interactions among CSP-A subunits in ARV and in rotavirus can explain the presence of internal D3 symmetry of the former and the absence of that symmetry in the latter. In ARV particles, N-terminal arms of the five ARV CSP-A subunits around a vertex anchor the polymerase and NTPase, while the arms of the five CSP-B subunits extend to interact with two other CSP-B arms around icosahedral threefold axes [8,9]. For the two oppositely directed threefolds that are the D3 symmetry axes, the CSP-B arm from one CSP decamer also contacts the RdRp at one of the threefold related vertices, thus determining the relative orientations of the corresponding assembly units. That is, the cyclic contacts between the arms of CSP-B subunits and RdRps generates threefold symmetric “supra-assemblies”, which become the north and south poles of the D3 cluster. The remaining six assembly units (one of which will not have an attached RNA segment and hence might lack a stably attached RdRp) could then fill in around the equator. The interactions that correlate the positions of the remaining five RdRps are not evident from published descriptions and coordinates; presumably these interactions are contacts (or exclusions) from other CSP-B arms.

The VP2 CSPs of rotaviruses have shorter N-terminal arms than those of the reoviruses, and they do not extend to contact an RdRp at a neighboring vertex. Thus, there is no mechanism to correlate the particular choice among five positions at one vertex with the choice at another. The second internal enzyme, VP3, contains all the cap forming activities, as rotaviruses do not have reovirus-like, external, capping-enzyme “turrets”. VP3 does not appear to have a fixed position with respect to the viral shell or with respect to the VP1 RdRp, and our reconstructions give no information about the interactions that recruit it to the assembling DLP. In the DLP_RNA reconstruction, there is an unassigned stretch of density associated with the RdRp N-terminal domain (Fig. S5c), but we cannot at the present resolution determine whether it could be a bound VP3 peptide or instead the N-terminal arm of a VP2 molecule, in addition to the two VP2A arms and one VP2B arm already assigned (Fig. 5).

### VP1 activation

The rotavirus RdRp is active *in vitro* only when associated with VP2, which is probably in the form of a decamer [22]. When particles assemble in the cytosol, association with both VP2 and the 3’ end of a (+) strand RNA segment incorporates VP1 and RNA into a precursor particle and enables VP1 to synthesize the (−) strand within that precursor. Addition of the TLP outer layer, when the DLP and VP4 enter the endoplasmic reticulum and acquire VP7, blocks further activity until loss of the VP4-VP7 outer layer releases the blockage, following infection of a new host cell. How does VP2 association activate (−) strand synthesis, and how does uncoating allow transcription -- i.e., (+) strand synthesis -- to start?

Comparison of the crystal structure of apo-VP1 with the VP1 structure in a non-transcribing particle (DLP or TLP) shows two distinct sets of conformational differences (Fig. 4). One set includes the flip of loop 261–271 enforced by association with VP2 and propagated in turn to the loop between residues 493 and 508. Analysis of the crystal structure of VP1 led to the suggestion that this loop might support the priming nucleotide, by analogy with its reovirus λ3 homolog [4]., Our new structures suggest that it senses binding with VP2 by transmitting the change at loop 261–271 to the vicinity of the catalytic site. The other set of conformational differences is displacement of the bracelet α-helical subdomain, residues 918–1006, also enforced by association with VP2 (to avoid collision with residues 348–362), resulting in constriction of template (or dsRNA) and transcript exit channels. The principal conformational changes accompanying addition of substrates to the DLP is likewise displacement of the α-helical subdomain, back to a position close to its position in the VP1 crystal structure; disordering of residues 346–373, in VP2A molecule 5, accommodates the shift and creates a channel for transcript release (Fig. 6 and 7).

How addition of the outer layer of the TLP suppresses the RdRp activity of the DLP is not as evident as the apparent mechanism for sensing association with VP2. The VP1-VP2 contacts appear to be the same in DLP and TLP, and our analysis of the structure does not indicate how inward displacement of VP2A tips (Fig. S7a), which accompanies addition of the outer layer, leads to inhibition of RdRp activity.

### Capping

Our reconstruction of a transcribing DLP shows that the likely transcript exit channel is continuous with an open passage to the pore on the fivefold (Fig. 7b). Density features we attribute to nascent mRNA suggest that transcripts do indeed pass directly from the catalytic site through the fivefold pore to the outside of the capsid. How, then, does the 5’ end of the transcript acquire a cap? A mechanism is in principle possible, in which the transcript emerges base-paired with the template from the template-exit channel, encounters VP3, and then separates and inserts into the transcript channel. Larger rearrangements in the C-terminal bracelet domain than those illustrated in Fig. 6 might allow such a shift, although none of our structures provide evidence for it. The relative positions of the NTPase and RdRp in ARV and CPV present a similar puzzle [10]. Priming of transcription by a VP3-produced, capped nucleotide, 3’-meGpppG-3’, or capped dinucleotide, 3’-meGpppGpG-3’, would provide a potential alternative mechanism for rotavirus. VP3 from disrupted DLPs appears indeed to synthesize these species [23]. The structure of VP3, recently determined but not yet published (BVV Prasad, personal communication), may help resolve this issue.

## Materials and Methods

### TLP and DLP purification

TLPs and DLPs were purified as previously described [25]. For TLP and DLP production, MA104 cells were grown in 850 cm^2^ roller bottles (Corning), and confluent monolayers were infected with rhesus rotavirus (RRV, G3 serotype) at MOI (multiplicity of infection) of 0.1 focus-forming unit (FFU)/cell in M199 medium supplemented with 1 mg/mL porcine pancreatic trypsin (Worthington Biochemical). Cell culture medium was collected 24–36 h post infection, when cell adherence was <5%. TLPs and DLPs were purified by freeze-thawing, ultracentrifugation, Freon-113 extraction, and separation on a cesium chloride gradient. TLPs were de-salted with a 5 ml Zeba spin column (Thermo Fisher) into 20 mM Tris pH 8.0, 100 mM NaCl, and 1 mM CaCl_2_ (TNC). DLPs were de-salted into 20 mM HEPES pH 7.5, 100 mM NaCl (HN).

For data collection of transcribing particles [19], 21 µl of TLPs (3.5 mg / ml) or DLPs (2.7 mg / ml) were added to a final reaction volume of 30 µl containing 150 mM NaCl; 9 mM MgCl_2_; 4 mM adenosine-5’-triphosphate (ATP) (New England Biolabs); 2 mM each of guanosine-5’-triphosphate (GTP), cytidine-5’-triphosphate (CTP), and uridine-5’-triphosphate (UTP) (New England Biolabs); and 640 µM S-adenosylmethionine (SAM) (New England Biolabs). The reaction was incubated for 5 min at 37 °C and placed on ice until grids were prepared.

### Data collection

A stack of pre-processed and selected 4178 DLP images was taken from a previously published dataset [26] (RRV, G3 serotype, 1024^2^ pixels, pixel size = 1.023 Å). For the other datasets (Table S1), we prepared cryo-EM grids by applying 3.2 µL of purified viral particles to glow-discharged C-flat grids (CF-1.2/1.3-4C, Protochips). We flash-cooled the grids after blotting off excess liquid by plunging them into liquid ethane, using a CP3 cryo plunger (Gatan). Blotting time was 4–6 s at about 88% relative humidity. On a Polara microscope operated at 300 kV and equipped with a K2 Summit detector (Gatan) we recorded with SerialEM (http://bio3d.colorado.edu/SerialEM) movies in super-resolution mode (40 frames, 0.2 s per frame, 8 electrons/Å^2^ per second) (Table S1).

### Image pre-processing

Movie frames were gain reference-corrected, aligned and dose-weighted with Motioncor2 (5×5 patch alignment, first two frames discarded) [27]. We Fourier-binned the aligned image sums to give a calibrated pixel size of 1.23 Å. We picked particles manually (TLP dataset) or detected them with Gautomatch (DLP_RNA, TLP_RNA). For calculation of template projections, done with EMAN2 [28] from a previous reconstruction, and automatic particle picking the angular sampling was 3° and the cutoff of the low-pass filter was 40 Å. We estimated defocus and astigmatism parameter values with Gctf using total-summed images and refinement for values at particle coordinates [29]. We extracted and normalized particles images with relion_preprocess [30] from the dose-weighted sums.

### Icosahedral reconstruction

We aligned the particles in FrealignX (Refine3D version 1.01, Reconstruct3D version 1.02) [31] using mode 3 (12 Å-resolution limit for alignment) and a low-pass filtered reference obtained from an atomic models (PDB 4v7q), followed by seven additional cycles in mode 1 (8–5 Å-resolution limit for alignment) (Fig. 2a). We applied icosahedral symmetry (setting I2) for projection matching and reconstruction and a spherical shell mask (TLP: inner radius = 210 Å, outer radius = 400 Å; DLP: inner radius = 210 Å, outer radius = 360 Å). Overall resolution estimates calculated from densities within a spherical shell containing VP2, VP6 and, in case of TLPs, VP7 are in Table S1.

### Local reconstruction of VP1 polymerase

For local reconstruction of VP1 polymerase, we used signal subtraction [11], sub-particle extraction [12], and classification methods [13]. Initially, we subtracted with relion_project [30] density from the original particle images, using a mask that included the entire viral particle, except all volume potentially occupied by VP1 (corresponding to PDB 4au6 that contains 5 overlapping copies of VP1 around the 5-fold axis) (Fig. 2b). A reconstruction calculated with icosahedral symmetry from this signal-subtracted particle stack showed, as expected, VP2, VP6 and, in case of TLPs, VP7 density erased.

We next extracted 60 sub-particles from each image of the original and the signal-subtracted particle stacks (Fig. 2c). Using Python (www.python.org) and the icosahedral alignment parameters (*φ_i_*, *θ_i_*, *ϕ_i_*, *x*_*i*_, *y*_*i*_) as input, we calculated extraction coordinates and sub-particle alignment parameters (*φ_s_*, *θ_s_*, *ϕ_s_*, *x*_*s*_, *y*_*s*_). Extraction coordinates are the projection, onto the *xy*-plane, of 60 points (initially obtained by applying the 60 icosahedral symmetry matrixes to a point laying on one of the 5-fold symmetry axis with a distance of 200 Å from the center of the virus, the approximate position of VP1) after rotation and translation according to the icosahedral alignment parameters (*φ_i_*, *θ_i_*, *ϕ_i_*, *x*_*i*_, *y*_*i*_). We used IMOD [32] to extract the sub-particles and create sub-particle stacks. Sub-particle alignment parameters (*φ*_*s*_, *θ*_*s*_, *ϕ*_*s*_, *x*_*s*_, *y*_*s*_) were calculated from the icosahedral alignment parameters (*φ*_*i*_, *θ*_*i*_, *ϕ*_*i*_, *x*_*i*_, *y*_*i*_) and the icosahedral symmetry matrixes such that insertion of the sub-particle image in Fourier space with C_1_ symmetry leads to a real space volume where all 60 icosahedral asymmetric subunits from a single virus are superimposed.

We used classification in FrealignX to determine which of the 5 possible positions the VP1 polymerase occupies on each vertex. For this we modified the FrealignX control script to allow class-specific masking of the reference volume for each of 5 classes (Fig. 2d). Refinement of alignment parameters was turned off during iterative classification (24–80 cycles). The resolution limit for classification was 5 Å. We also effectively turned off per-particle weighting in the reconstruction step (BSC = 0.0). As theoretically expected, we obtained 5 classes with approximately equal particle numbers that are related by a 5-fold rotational symmetry axis). Final reconstructions were obtained by applying the result of the classification step to the original, non-signal-subtracted subparticle stack (Fig. 2e). Fourier shell correlation (FSC) plots of the local reconstructions are in Fig. S1.

### Model building and refinement

We placed the VP1 crystal structure (PDB 2R7Q) and ten copies of VP2 (PDB 4V7Q) into the DLP density map with PyMOL (The PyMOL Molecular Graphics System, Version 2.1 Schrödinger, LLC). We manually adjusted and completed the rigid body-fitted subunits and also built the VP2 N termini in contact with VP1 using the program O [33]. Atomic coordinates and *B*-factors were refined with PHENIX (phenix.real_space_refine) [34]. In addition to standard stereochemical restraints and *B*-factor restraints, we used Ramachandran, rotamer, secondary structure restraints, and reference model restraints (Table S2). Residues included in the models are summarized in Table S2. We validated the models with MolProbity [35]. Model statistics are in Table S1 and Fig. S1.

### Figure preparation

We prepared the figures with PyMOL (The PyMOL Molecular Graphics System, Version 2.1 Schrödinger, LLC), POV-Ray (www.povray.org) and matplotlib [36].

### Accession numbers for atomic coordinates and density maps

The cryo-EM local reconstructions are deposited in the Electron Microscopy Data Bank (EMDB IDs: EMD-20086, EMD-20087, EMD-20088, EMD-20089). The atomic coordinates of the refined models are deposited in the Protein Data Bank (PDB IDs: 6OJ3, 6OJ4, 6OJ5, 6OJ6).

## Authorship contribution statement

**Simon Jenni:** Conceptualization; Data curation; Formal analysis; Investigation; Methodology; Validation; Writing - original draft; Writing - review & editing. **Eric N. Salgado:** Investigation; Methodology; Resources. **Tobias Herrmann:** Data curation; Formal analysis; Investigation. **Zongli Li:** Data curation; Investigation. **Timothy Grant:** Data curation; Investigation; Methodology; Resources; Software. **Nikolaus Grigorieff:** Data curation; Investigation; Methodology; Resources; Software. **Stefano Trapani:** Conceptualization; Formal analysis; Methodology. **Leandro F. Estrozi:** Conceptualization; Formal analysis; Methodology. **Stephen C. Harrison:** Conceptualization; Formal analysis; Funding acquisition; Investigation; Methodology; Project administration; Supervision; Validation; Writing - original draft; Writing - review & editing.

## Acknowledgments

We thank Adam Johnson and Stephen Hinshaw for help with EM data collection and acknowledge support from NIH grant CA-13202 (to SCH). SCH is an Investigator in the Howard Hughes Medical Institute.

## Abbreviations

3’CS+: positive strand 3’ consensus sequence
ARV: aquareoviurs
CPV: cytoplasmic polyhedrosis virus
CSP: capsid-shell protein
DLP: double-layer particle
dsRNA: double-stranded RNA
FFU: focus-forming unit
FSC: Fourier shell correlation
MOI: multiplicity of infection
NTP: nucleotide triphosphate
NTPase: nucleoside triphosphatase
RdRp: RNA-dependent RNA polymerase
RRV: rhesus rotavirus
SAM: S-adenosyl methionine
TLP: triple-layer particle

---

**Fig. S1.**
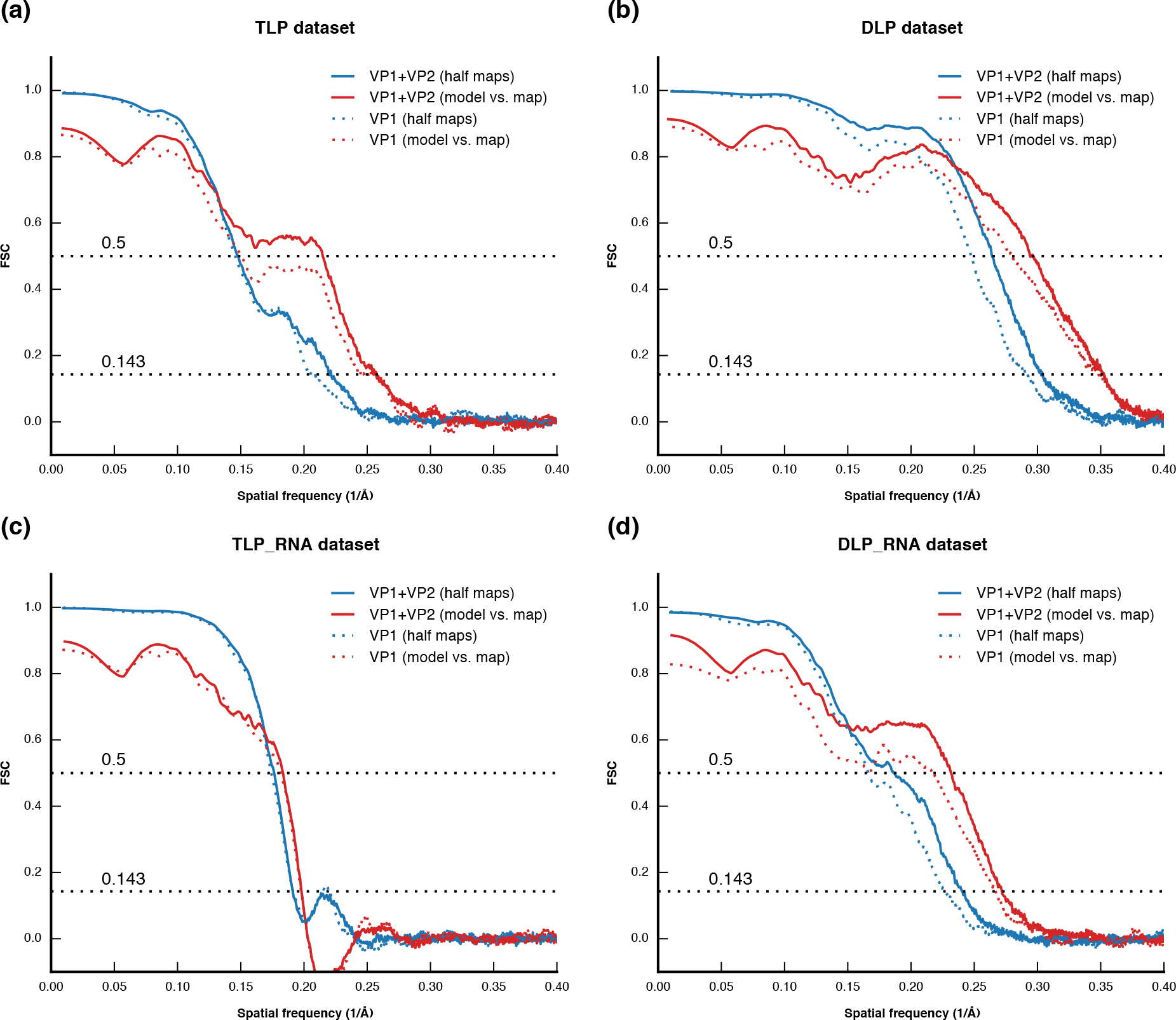
Fourier shell correlation (FSC) between the two half-maps (blue), and between the final map and the refined model, calculated with phenix.mtriage [37]. Solid lines are after applying a mask encompassing VP1 and VP2, dash lines after masking VP1 only. (a) “Triple-layer particle” (TLP) reconstruction. (b) “Double-layer particle” (DLP) reconstruction. (c) TLP_RNA reconstruction after incubation with nucleotides and S-adenosyl methionine (SAM). (d) DLP_RNA reconstruction after incubation with nucleotides and SAM.

**Fig. S2.**
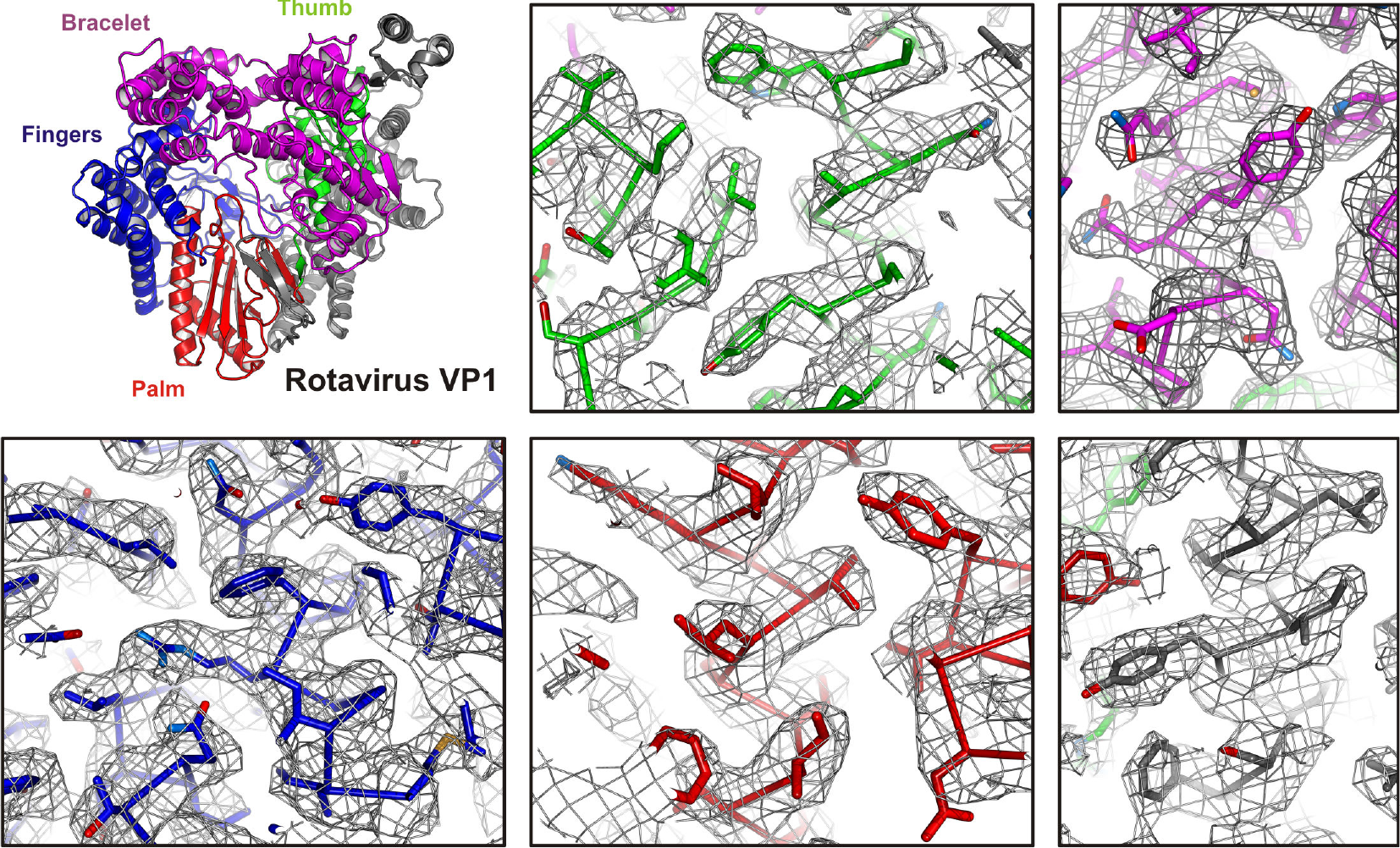
Representative regions of VP1 in the DLP cryo-EM map (EMD-20087). An overview of the VP1 structure is shown (top left) with its domains colored: N-terminal domain, gray; palm, red; thumb, green; fingers blue; bracelet domain, magenta. Density is shown as gray mesh.

**Fig. S3.**
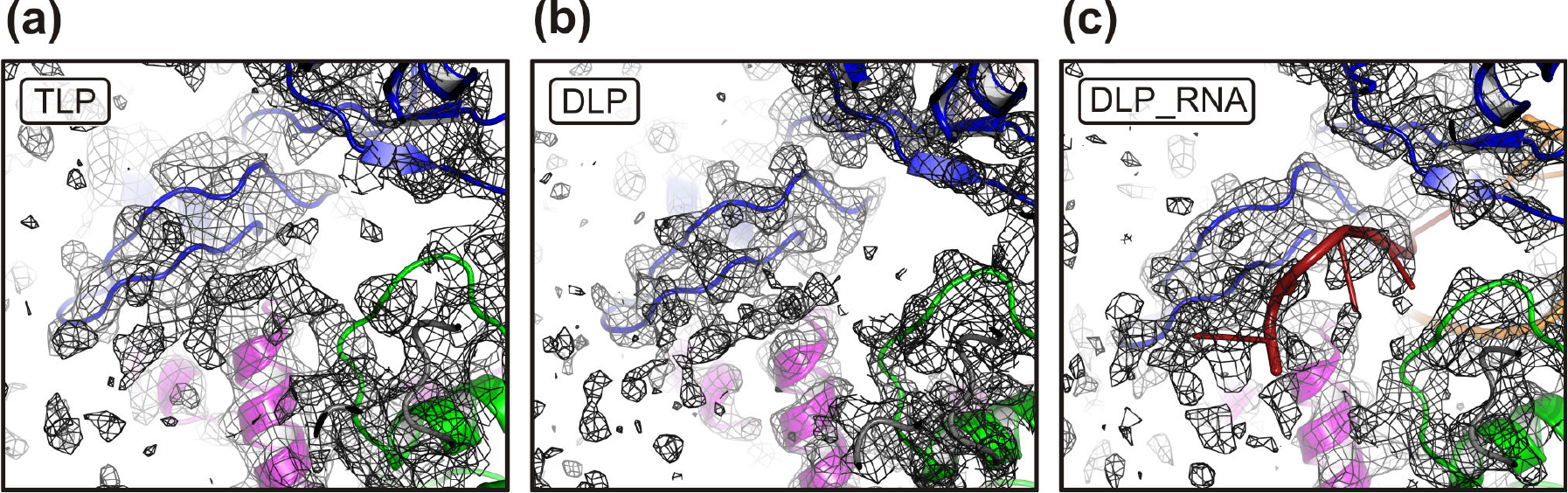
Close-up views of the RdRp template-entry tunnel. The polymerase is colored as in Fig. 3. Cryo-EM density is shown as black mesh. (a) TLP reconstruction. (b) DLP reconstruction. (c) TLP_RNA reconstruction after incubation with nucleotides. RNA template and transcript strands are in dark red and orange, respectively.

**Fig. S4.**
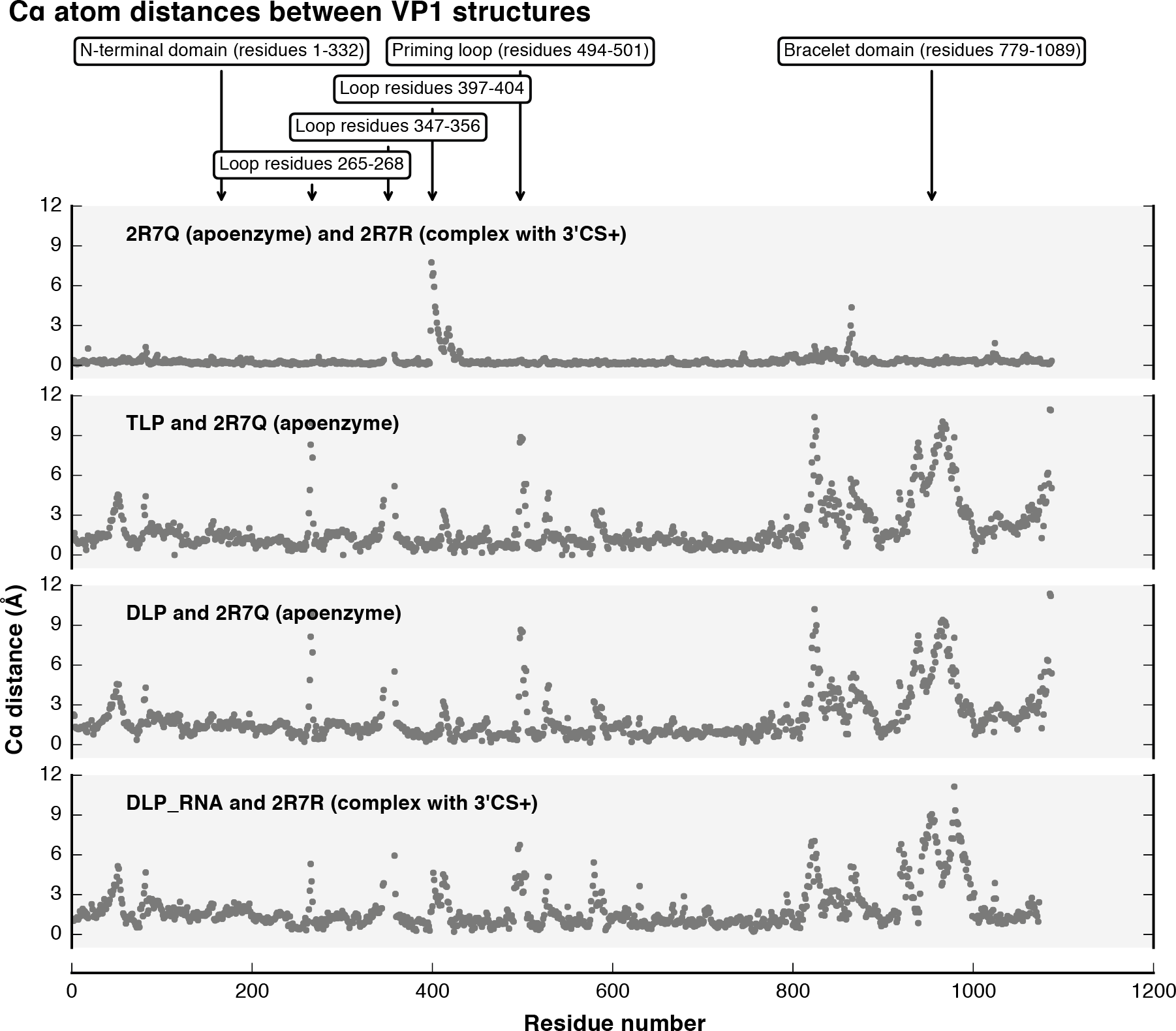
Structural differences between VP2-bound RdRp and the crystal structures obtained in isolation [4]. Structures were superimposed using residues of the fingers, palm and thumb domains only. The plots show the distances between corresponding Cα atoms after superposition. Regions with large conformational shifts are labeled on top.

**Fig. S5.**
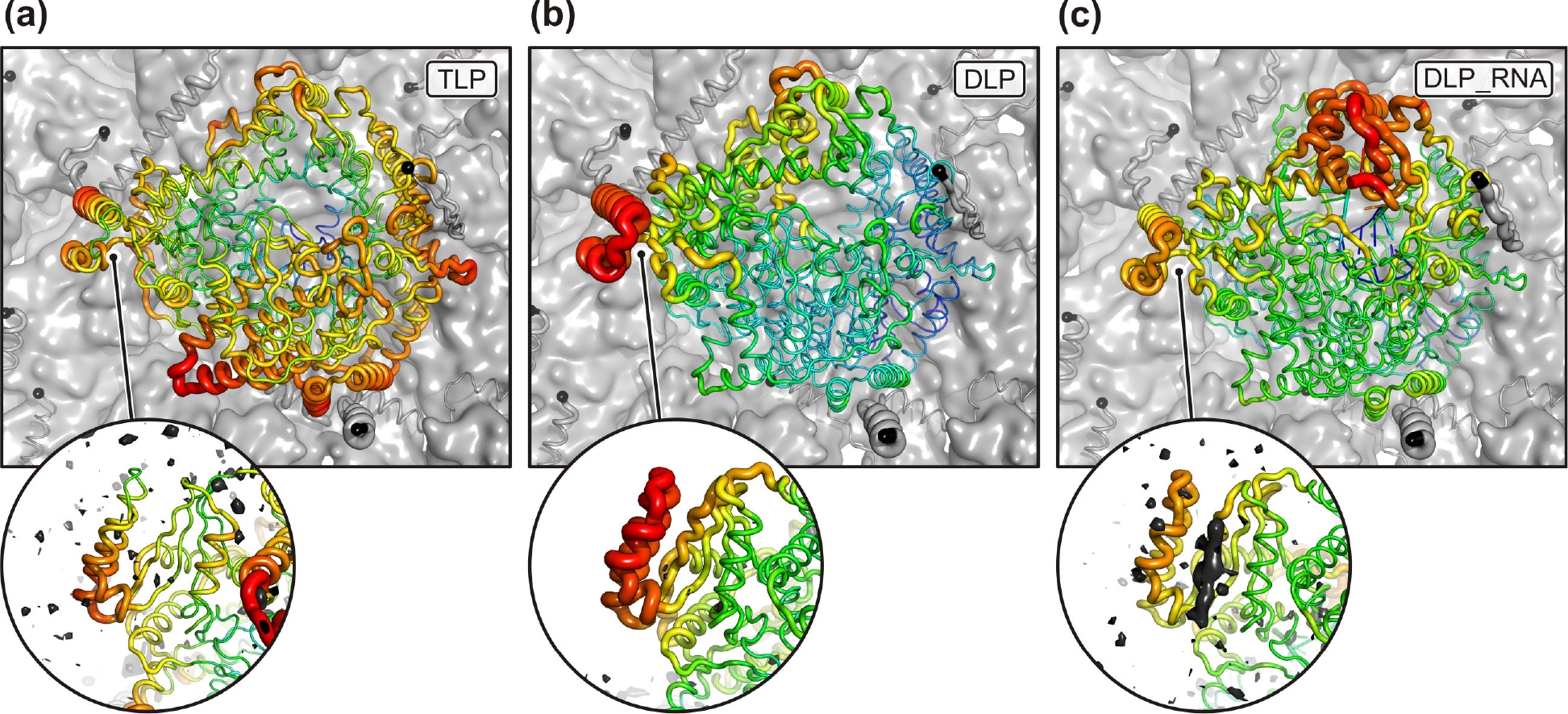
Temperature factors of VP2-bound RdRp. Magnified insets show the N-terminal RdRp domain with a difference map (cryo-EM map - model) shown in black and calculated to 4.5 Å resolution. Because of the different overall temperature factors of the three reconstructions, temperature factors cannot be compared across the panels and are only indicative of relative differences within one structure. (a) TLP reconstruction. (b) DLP reconstruction. (c) DLP_RNA reconstruction after incubation with nucleotides. Additional density bound to the N-terminal RdRp domain is visible DLP_RNA reconstruction.

**Fig. S6.**
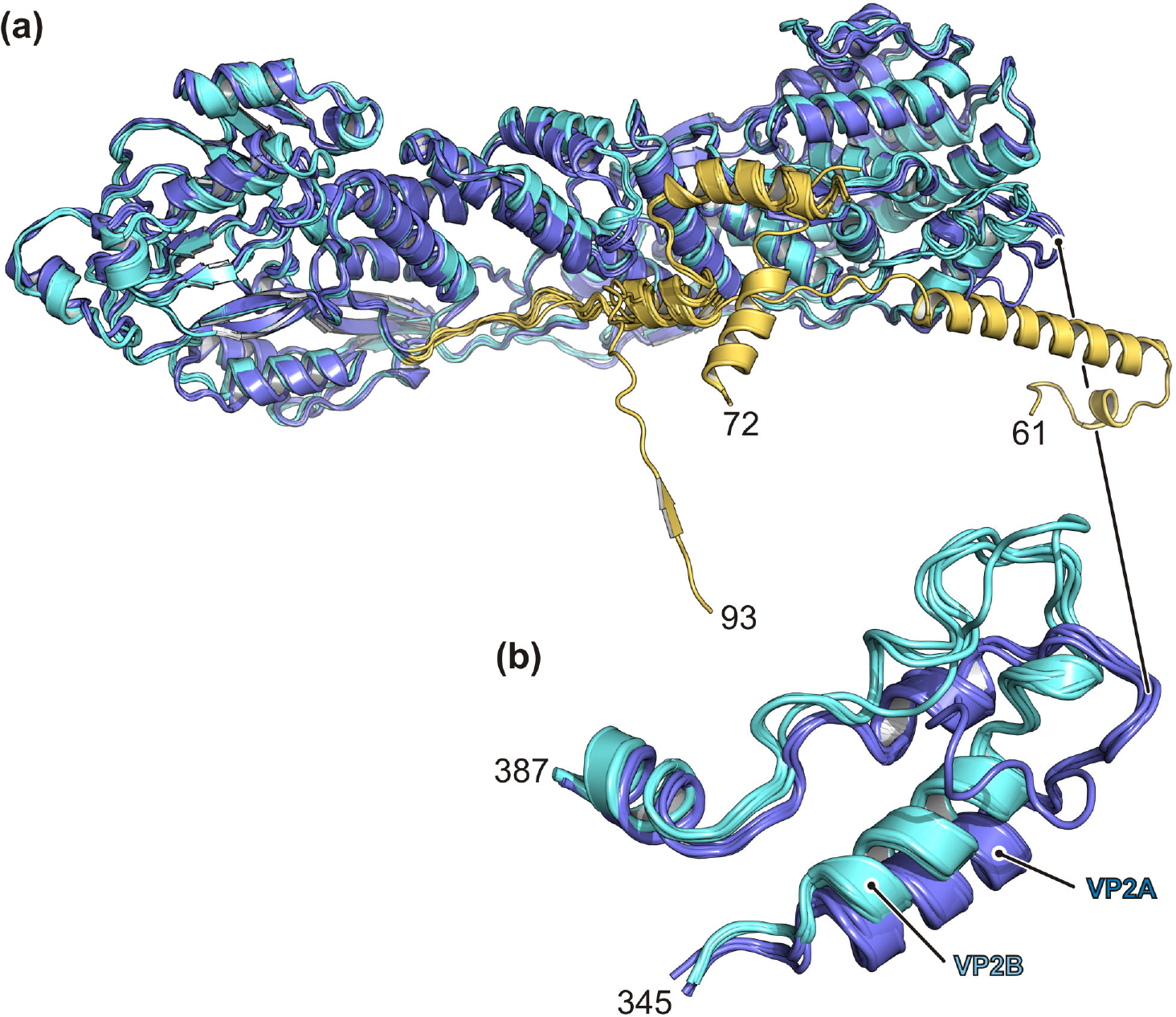
Structural comparison of the VP2 molecules after superimposing residues 124–887 (refined model from DLP dataset). In the ribbon representation VP2A is colored in blue, and VP2B in cyan, except for the N-terminal residues, which are colored in yellow. Residue numbers are indicated. (a) Overview showing conformational variability of the N termini. (b) Magnification of the VP2 tip.

**Fig. S7.**
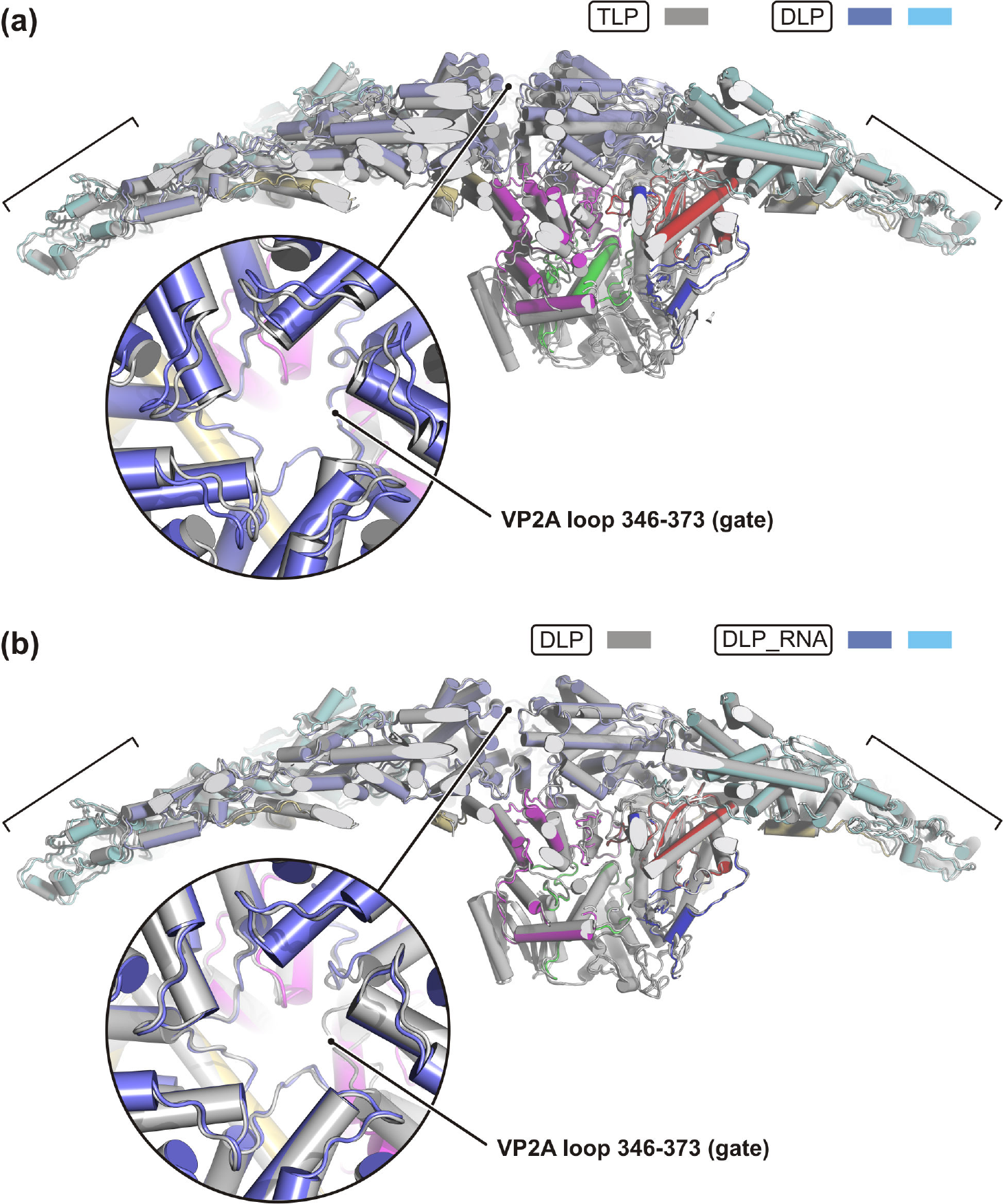
Structural comparison of the local reconstructions. The side views are partially cut and show the VP2 decamers with bound VP1 after superposing VP2B peripheral residues (124–268, 682–851), as indicated by brackets. The insets show a magnified view along the icosahedral 5-fold axis from the outside, after superposing VP2A proximal residues (321–602). Tips of the VP2A subunits surround the narrowest part of the capsid pore. (a) Comparison of the TLP (gray) and DLP (colored) structures. (b) Comparison of the DLP (gray) and DLP_RNA (colored) structures. The template and transcript strands are omitted in the figure for clarity.

**Fig. S8.**
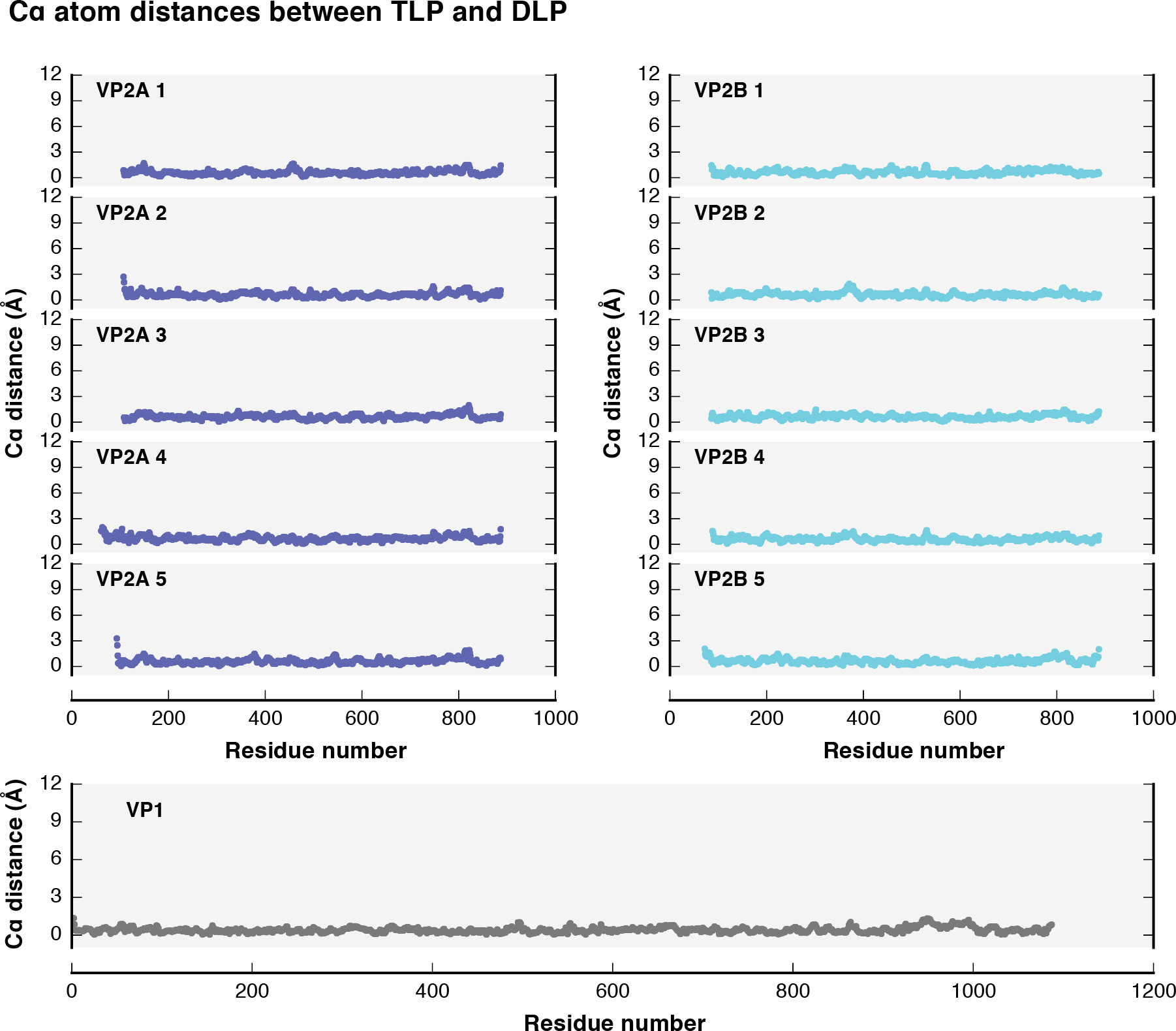
Structural differences between TLP and DLP. Subunits were superposed individually and the plots show the distances between corresponding Cα atoms after superposition.

**Fig. S9.**
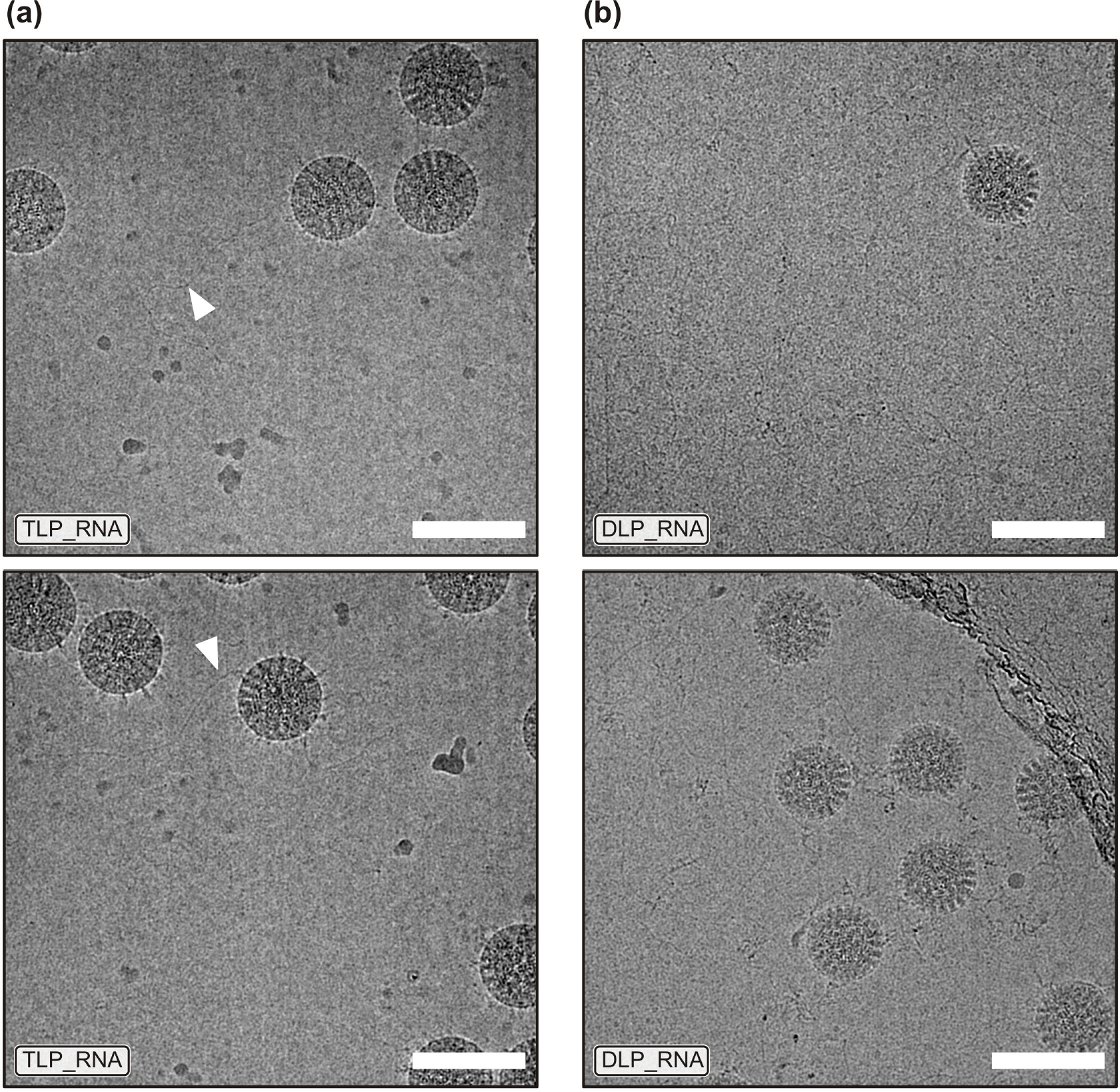
Cryo-EM images of rotavirus particles in vitreous ice after incubation with nucleotides. The images shown are low-pass filtered at 15 Å resolution. The scale bar is 100 nm. (a) TLPs only occasionally showed RNA strands emerging from viruses (indicated by arrowheads) and the RdRp reconstruction did not show any RNA density in the active site. (b) DLPs with emerging RNA strands. Released transcripts are also visible in the background.

**Fig. S10.**
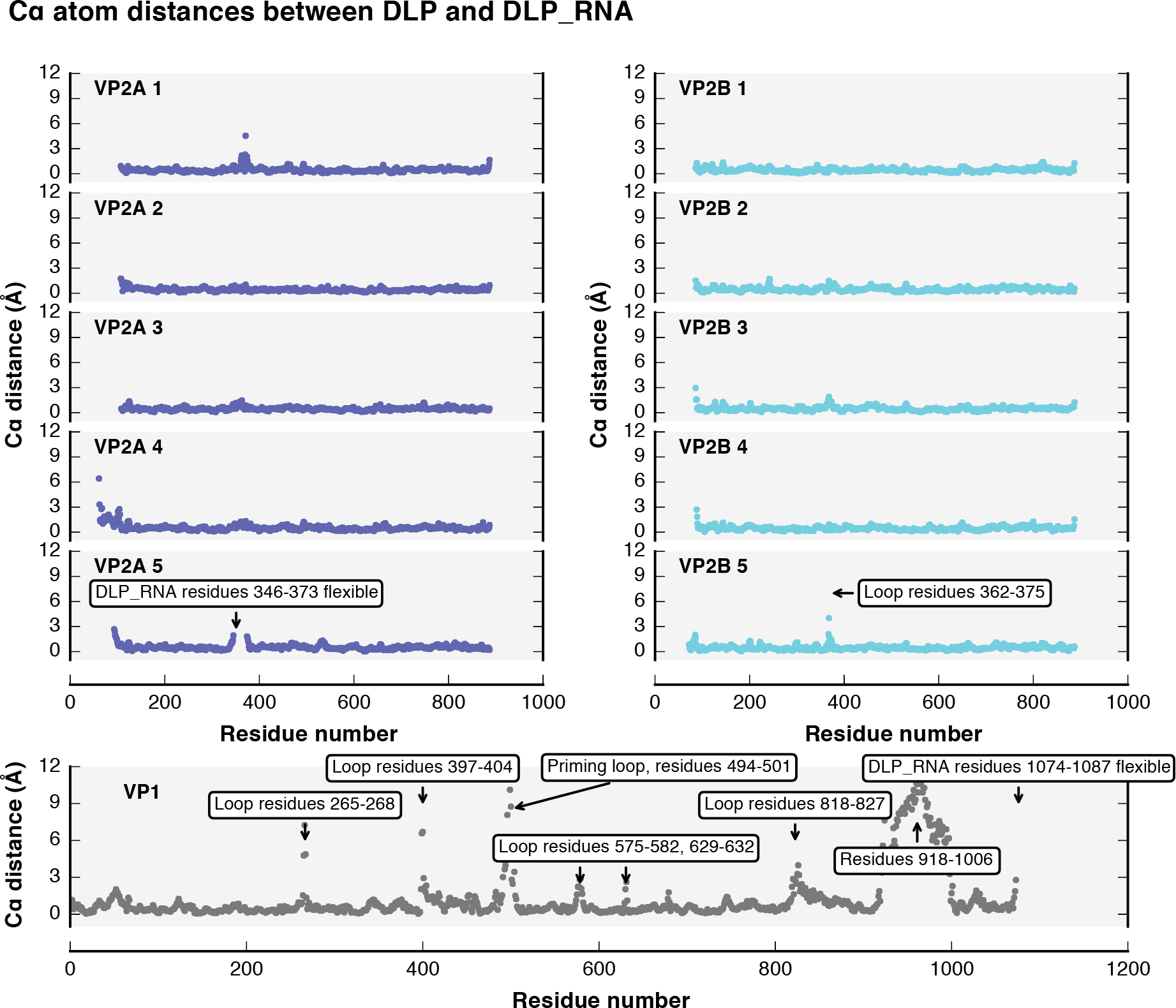
Structural differences between DLP and DLP_RNA. Subunits were superposed individually and the plots show the distances between corresponding Cα atoms after superposition.

**Fig. S11.**
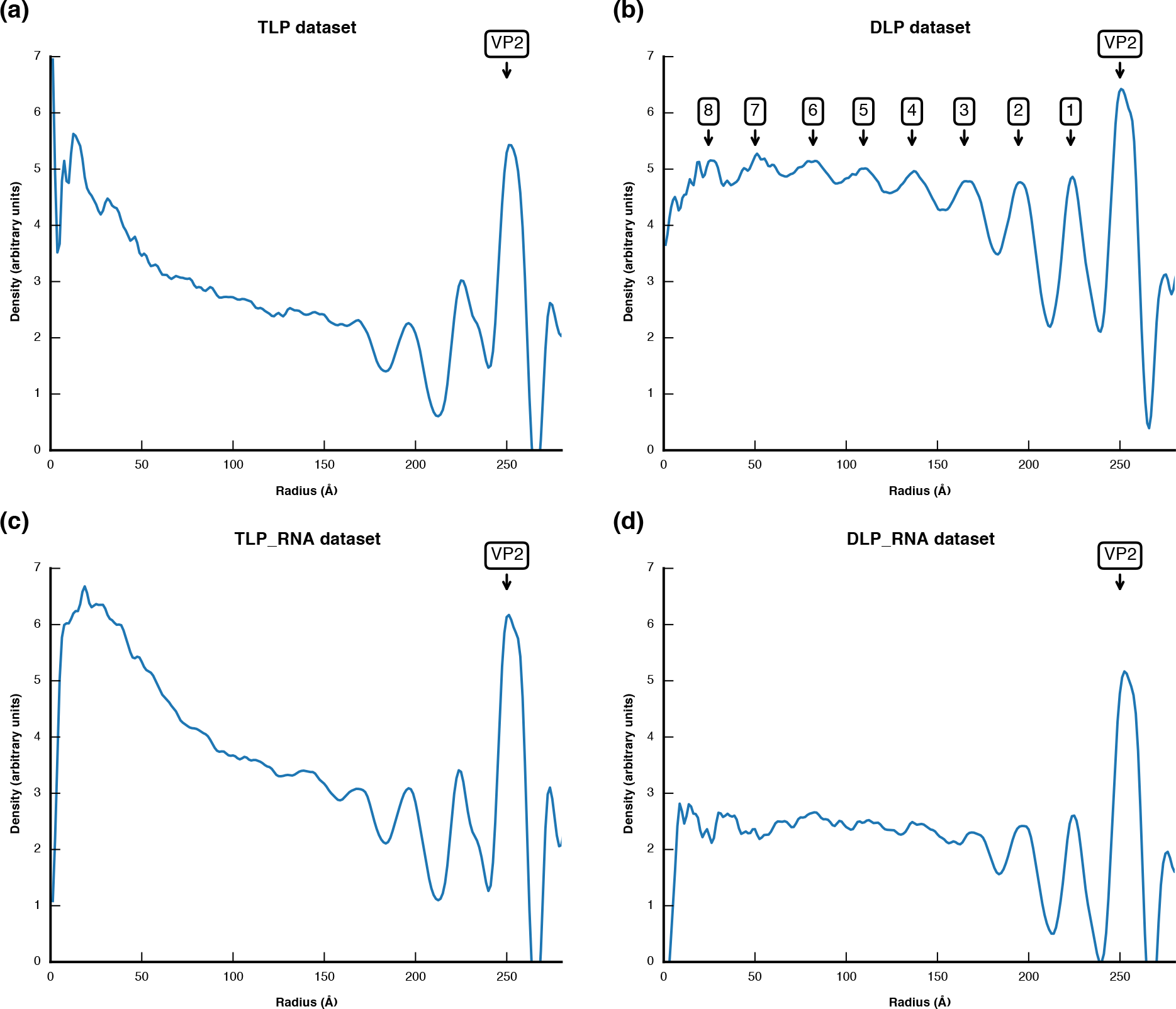
Spherical density average of the icosahedral reconstructions. The peak corresponding to the VP2 capsid is labeled. (a) TLP reconstruction. (b) DLP reconstruction. Peaks corresponding to spherical layers of dsRNA are sequentially numbered. (c) TLP_RNA reconstruction after incubation with nucleotides. (b) DLP_RNA reconstruction after incubation with nucleotides.

**Fig. S12.**
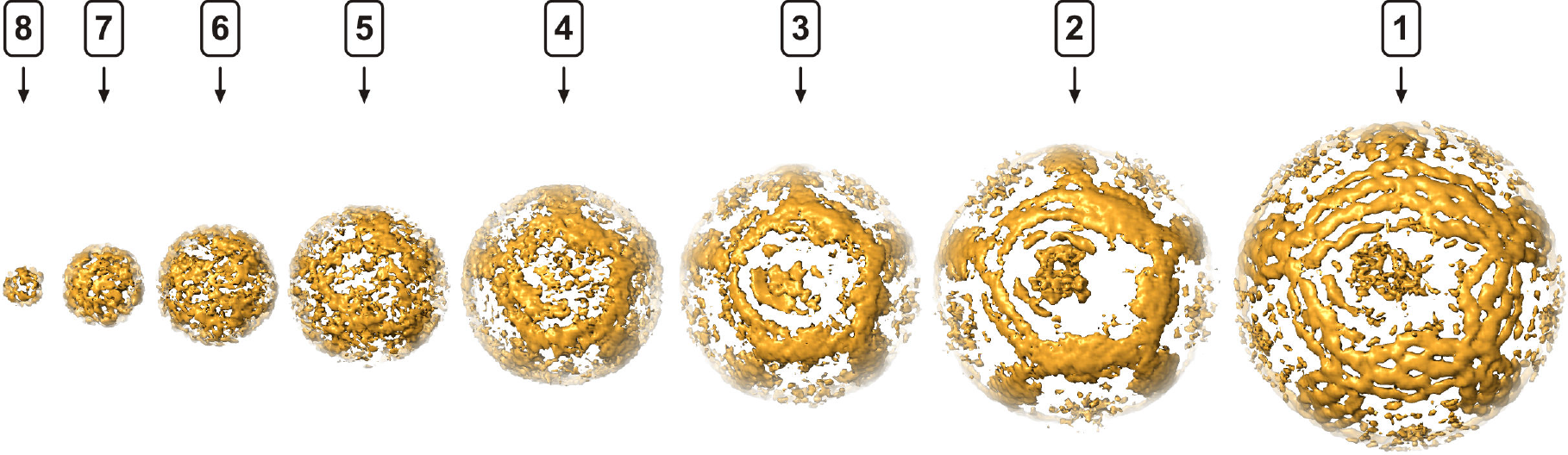
dsRNA genome density (DLP dataset). The concentric layers containing most of dsRNA density (Fig. S11) are masked and shown individually. The view is along the fivefold symmetry axis where VP1 is aligned.

**Fig. S13.**
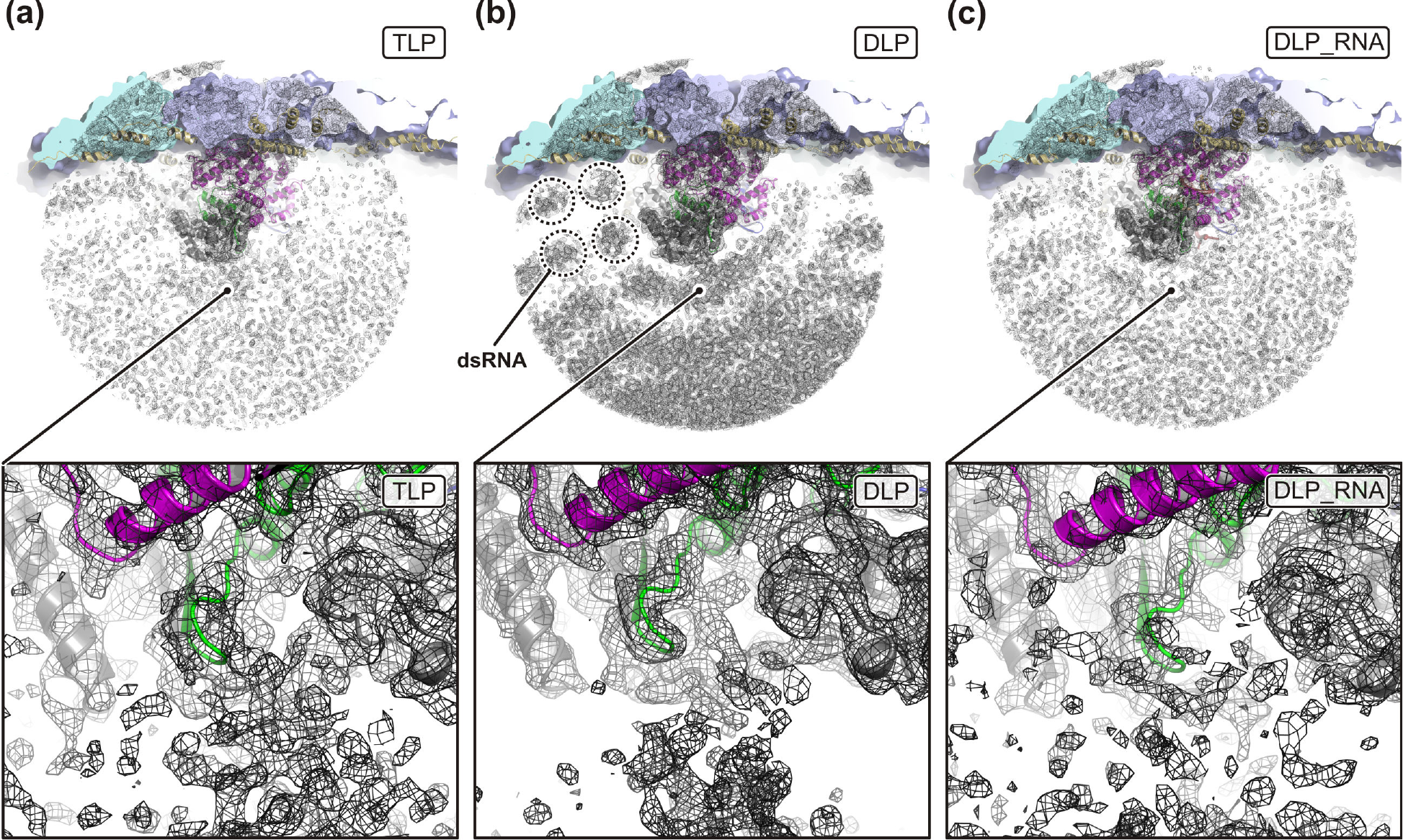
RdRp cap binding site. Structures are colored as in Fig. 3. Cryo-EM density is shown as black mesh. The close-up views focus on the cap-binding site, located between the RdRp N-terminal (gray) and thumb (green) domains. (a) TLP reconstruction. (b) DLP reconstruction. Dashed circles indicate density of dsRNA segments, approximately cross-sectioned. (c) TLP_RNA reconstruction after incubation with nucleotides.

**Table S1.** Cryo-EM data collection and model statistics.

|  | TLP | DLP | TLP_RNA | DLP_RNA |
| --- | --- | --- | --- | --- |
|  | EMD-20086<br>PDB-6OJ3 | EMD-20087<br>PDB-6OJ4 | EMD-20088<br>PDB-6OJ5 | EMD-20089<br>PDB-6OJ6 |
|  | RRV<br>Serotype G3 | RRV<br>Serotype G3 | RRV<br>Serotype G3 | RRV<br>Serotype G3 |
| **Data collection** |  |  |  |  |
| Electron microscope | Polara | Krios | Polara | Polara |
| Magnification | 40650 | 48876 | 40650 | 40650 |
| Voltage (kV) | 300 | 300 | 300 | 300 |
| Defocus range ($\mu\text{m}$)<sup>a</sup> | 0.5–2.5 | 0.5–1.8 | 0.5–2.5 | 0.4–1.5 |
| Pixel size ($\text{\AA}$) | 1.23 | 1.023 | 1.23 | 1.23 |
| **Icosahedral reconstruction** |  |  |  |  |
| Number of images | 2547 | 4178 | 6822 | 1607 |
| Box size (pixels) | 1024 | 1024 | 1024 | 840 |
| Symmetry imposed | I (setting I2) | I (setting I2) | I (setting I2) | I (setting I2) |
| Map resolution ($\text{\AA}$)<sup>b</sup> | 4.2 | 3.2 | 4.2 | 4.0 |
| **Local reconstruction** |  |  |  |  |
| Number of images | 30606 | 50176 | 82008 | 19321 |
| Box size (pixels)<sup>c</sup> | 300 (264) | 352 (320) | 300 (264) | 300 (264) |
| Symmetry imposed | C<sub>1</sub> | C<sub>1</sub> | C<sub>1</sub> | C<sub>1</sub> |
| Map resolution ($\text{\AA}$)<sup>d</sup> | 4.5 | 3.3 | 5.2 | 4.2 |
| Sharpening *B* factor ($\text{\AA}^2$) | -60.0 | -40.0 | -60.0 | -60.0 |
| **Model statistics** |  |  |  |  |
| Refinement resolution ($\text{\AA}$) | 4.5 | 3.2 | 4.2 | 4.0 |
| CC (mask) | 0.74 | 0.79 | 0.78 | 0.72 |
| Model composition |  |  |  |  |
| Non-hydrogen atoms | 74075 | 74095 | 74095 | 74265 |
| Protein residues | 9072 | 9072 | 9072 | 9030 |
| Nucleotides | 0 | 0 | 0 | 24 |
| *B* factors ($\text{\AA}^2$) | | | | |
| Protein | 90.5 | 48.8 | 193.0 | 66.7 |
| RNA | N.A. | N.A. | N.A. | 97.8 |
| R.m.s. deviations |  |  |  |  |
| Bond lengths ($\text{\AA}$) | 0.008 | 0.009 | 0.008 | 0.008 |
| Bond angles ($^\circ$) | 1.290 | 0.868 | 1.391 | 1.256 |
| Validation |  |  |  |  |
| MolProbity clash score | 3.5 | 3.9 | 4.2 | 3.1 |
| Poor rotamers (%) | 0.0 | 0.0 | 0.0 | 0.0 |
| Ramachandran plot |  |  |  |  |
| Favored (%) | 91.0 | 90.7 | 90.8 | 90.7 |
| Allowed (%) | 9.0 | 9.3 | 9.2 | 9.3 |
| Disallowed (%) | 0.0 | 0.0 | 0.1 | 0.1 |

<sup>a</sup> Approximate range of underfocus<sup>b</sup> Resolution where Fourier shell correlation (FSC) between half-maps drops below 0.143 after applying a spherical shell mask (TLP: inner radius = 210 $\text{\AA}$ , outer radius = 400 $\text{\AA}$ ; DLP: inner radius = 210 $\text{\AA}$ , outer radius = 360 $\text{\AA}$ )<sup>c</sup> Pixel number in parenthesis used during intermediate processing steps<sup>d</sup> Resolution where FSC between half-maps drops below 0.143 after applying a mask encompassing VP1 and 10 copies of VP2

**Table S2.** Model composition.

|  | **TLP** | **DLP** | **TLP_RNA** | **DLP_RNA** |
| --- | --- | --- | --- | --- |
|  | EMD-20086<br>PDB-6OJ3 | EMD-20087<br>PDB-6OJ4 | EMD-20088<br>PDB-6OJ5 | EMD-20089<br>PDB-6OJ6 |
|  | RRV <sup>a</sup><br>Serotype G3 | RRV <sup>a</sup><br>Serotype G3 | RRV <sup>a</sup><br>Serotype G3 | RRV <sup>a</sup><br>Serotype G3 |
| **Residues modeled** |  |  |  |  |
| VP2A 1 (chain A) | 107–887 | 107–887 | 107–887 | 107–887 |
| VP2A 2 (chain B) | 107–887 | 107–887 | 107–887 | 107–887 |
| VP2A 3 (chain C) | 108–887 | 108–887 | 108–887 | 108–887 |
| VP2A 4 (chain D) | 61–887 | 61–887 | 61–887 | 61–887 |
| VP2A 5 (chain E) | 93–887 | 93–887 | 93–887 | 93–345,<br>374–887 |
| VP2B 1 (chain F) | 86–887 | 86–887 | 86–887 | 86–887 |
| VP2B 2 (chain G) | 86–887 | 86–887 | 86–887 | 86–887 |
| VP2B 3 (chain H) | 86–887 | 86–887 | 86–887 | 86–887 |
| VP2B 4 (chain I) | 88–887 | 88–887 | 88–887 | 88–887 |
| VP2B 5 (chain J) | 72–887 | 72–887 | 72–887 | 72–887 |
| VP1 (chain P) | 2–1087 | 2–1087 | 2–1087 | 2–1073 |
| -RNA template (chain T) | N.A. | N.A. | N.A. | 97–110 |
| +RNA transcript (chain U) | N.A. | N.A. | N.A. | 101–110 |
| **NCS restraints** |  |  |  |  |
| Residue range | (A,B,C,D,E)<br>125–360,<br>373–887<br>(F,G,H,I,J)<br>95–887 | (A,B,C,D,E)<br>125–360,<br>373–887<br>(F,G,H,I,J)<br>95–887 | (A,B,C,D,E)<br>125–360,<br>373–887<br>(F,G,H,I,J)<br>95–887 | (A,B,C,D,E)<br>125–360,<br>373–887<br>(F,G,H,I,J)<br>95–887 |
| **Reference model restraints** |  |  |  |  |
| Reference model | DLP | PDB-2R7R | DLP | DLP |
| Residue range | (A,B,C,D,E,<br>F,G,H,I) all<br>(J) all <sup>b</sup><br>(P) all <sup>c</sup> | (P) 27–71 | (A,B,C,D,E,<br>F,G,H,I) all<br>(J) all <sup>b</sup><br>(P) all <sup>c</sup> | (A,B,C,D,E,<br>F,G,H,I) all<br>(J) all <sup>b</sup><br>(P) all <sup>c</sup> |

<sup>a</sup> GenBank accession numbers: VP2, EU636925; VP1, EU636924.
<sup>b</sup> except residues 357–377
<sup>c</sup> except residues 263–269, 490–505, 920–925

**Table S3.** Statistical analysis of V1 binding on DLP fivefold axes ^a^.

|  | Position 1 | Position 2 | Position 3 | Position 4 | Position 5 |
| --- | --- | --- | --- | --- | --- |
| Fivefold 2 | 159 | 188 | 160 | 170 | 172 |
| Fivefold 3 | 157 | 195 | 177 | 140 | 178 |
| Fivefold 4 | 167 | 163 | 176 | 167 | 175 |
| Fivefold 5 | 134 | 168 | 195 | 193 | 160 |
| Fivefold 6 | 175 | 182 | 159 | 177 | 156 |
| Fivefold 7 | 167 | 176 | 174 | 170 | 163 |
| Fivefold 8 | 186 | 161 | 177 | 170 | 156 |
| Fivefold 9 | 144 | 186 | 165 | 162 | 168 |
| Fivefold 10 | 171 | 166 | 147 | 186 | 175 |
| Fivefold 11 | 169 | 149 | 175 | 165 | 189 |
| Fivefold 12 | 144 | 173 | 162 | 202 | 168 |

<sup>a</sup> For 848 particles (20.3% of the DLP dataset), where V1 occupies position 1 on the fivefold 1 axis, we counted the number of VP1 occupying position 1, 2, 3, 4 or 5 on each other fivefold axis.

## References

[1] Estes MK, Greenberg H. Rotaviruses. In: Knipe DM, Howley PM, editors. Fields Virology. Philadelphia, PA: Lippincott Williams and Wilkins; 2013. p. 1347–401.

[2] Trask SD, Ogden KM, Patton JT. Interactions among capsid proteins orchestrate rotavirus particle functions. Curr. Opin. Virol. 2012;2:373–9.

[3] McClain B, Settembre E, Temple BR, Bellamy AR, Harrison SC. X-ray crystal structure of the rotavirus inner capsid particle at 3.8 A resolution. J. Mol. Biol. 2010;397:587–99.

[4] Lu X, McDonald SM, Tortorici MA, Tao YJ, Vasquez-Del Carpio R, Nibert ML, et al. Mechanism for coordinated RNA packaging and genome replication by rotavirus polymerase VP1. Structure. 2008;16:1678–88.

[5] Tao Y, Farsetta DL, Nibert ML, Harrison SC. RNA synthesis in a cage--structural studies of reovirus polymerase lambda3. Cell. 2002;111:733–45.

[6] McDonald SM, Tao YJ, Patton JT. The ins and outs of four-tunneled Reoviridae RNA- dependent RNA polymerases. Curr. Opin. Struct. Biol. 2009;19:775–82.

[7] Estrozi LF, Settembre EC, Goret G, McClain B, Zhang X, Chen JZ, et al. Location of the dsRNA-dependent polymerase, VP1, in rotavirus particles. J. Mol. Biol. 2013;425:124–32.

[8] Wang X, Zhang F, Su R, Li X, Chen W, Chen Q, et al. Structure of RNA polymerase complex and genome within a dsRNA virus provides insights into the mechanisms of transcription and assembly. Proc. Natl. Acad. Sci. U. S. A. 2018;115:7344–9.

[9] Ding K, Nguyen L, Zhou ZH. In Situ Structures of the Polymerase Complex and RNA Genome Show How Aquareovirus Transcription Machineries Respond to Uncoating. J. Virol. 2018;92.

[10] Li X, Zhou N, Chen W, Zhu B, Wang X, Xu B, et al. Near-Atomic Resolution Structure Determination of a Cypovirus Capsid and Polymerase Complex Using Cryo-EM at 200kV. J. Mol. Biol. 2017;429:79–87.

[11] Bai XC, Rajendra E, Yang G, Shi Y, Scheres SH. Sampling the conformational space of the catalytic subunit of human gamma-secretase. Elife. 2015;4.

[12] Ilca SL, Kotecha A, Sun X, Poranen MM, Stuart DI, Huiskonen JT. Localized reconstruction of subunits from electron cryomicroscopy images of macromolecular complexes. Nat. Commun. 2015;6:8843.

[13] Lyumkis D, Brilot AF, Theobald DL, Grigorieff N. Likelihood-based classification of cryo-EM images using FREALIGN. J. Struct. Biol. 2013;183:377–88.

[14] McDonald SM, Patton JT. Assortment and packaging of the segmented rotavirus genome. Trends Microbiol. 2011;19:136–44.

[15] Zeng CQ, Estes MK, Charpilienne A, Cohen J. The N terminus of rotavirus VP2 is necessary for encapsidation of VP1 and VP3. J. Virol. 1998;72:201–8.

[16] Salgado EN, Upadhyayula S, Harrison SC. Single-particle detection of transcription following rotavirus entry. J. Virol. 2017;91:e00651–17.

[17] Lawton JA, Estes MK, Prasad BV. Comparative structural analysis of transcriptionally competent and incompetent rotavirus-antibody complexes. Proc. Natl. Acad. Sci. U. S. A. 1999;96:5428–33.

[18] Chen JZ, Settembre EC, Aoki ST, Zhang X, Bellamy AR, Dormitzer PR, et al. Molecular interactions in rotavirus assembly and uncoating seen by high-resolution cryo-EM. Proc. Natl. Acad. Sci. U. S. A. 2009;106:10644–8.

[19] Lawton JA, Estes MK, Prasad BV. Three-dimensional visualization of mRNA release from actively transcribing rotavirus particles. Nat. Struct. Biol. 1997;4:118–21.

[20] Settembre EC, Chen JZ, Dormitzer PR, Grigorieff N, Harrison SC. Atomic model of an infectious rotavirus particle. EMBO J. 2011;30:408–16.

[21] Rickgauer JP, Grigorieff N, Denk W. Single-protein detection in crowded molecular environments in cryo-EM images. Elife. 2017;6.

[22] Gallegos CO, Patton JT. Characterization of rotavirus replication intermediates: a model for the assembly of single-shelled particles. Virology. 1989;172:616–27.

[23] Chen D, Patton JT. De novo synthesis of minus strand RNA by the rotavirus RNA polymerase in a cell-free system involves a novel mechanism of initiation. RNA. 2000;6:1455–67.

[24] Ding K, Celma C, Zhang X, Chang T, Shen W, Atanasov I, et al. In situ structures of a dsRNA virus polymerase in action and mechanism of mRNA transcription and release. Nature Communications, submitted. 2019.

[25] Aoki ST, Settembre EC, Trask SD, Greenberg HB, Harrison SC, Dormitzer PR. Structure of rotavirus outer-layer protein VP7 bound with a neutralizing Fab. Science. 2009;324:1444–7.

[26] Grant T, Grigorieff N. Measuring the optimal exposure for single particle cryo-EM using a 2.6 A reconstruction of rotavirus VP6. Elife. 2015;4:e06980.

[27] Zheng SQ, Palovcak E, Armache JP, Verba KA, Cheng Y, Agard DA. MotionCor2: anisotropic correction of beam-induced motion for improved cryo-electron microscopy. Nat. Methods. 2017;14:331–2.

[28] Bell JM, Chen M, Baldwin PR, Ludtke SJ. High resolution single particle refinement in EMAN2.1. Methods. 2016;100:25–34.

[29] Zhang K. Gctf: Real-time CTF determination and correction. J. Struct. Biol. 2016;193:1–12.

[30] Scheres SH. RELION: implementation of a Bayesian approach to cryo-EM structure determination. J. Struct. Biol. 2012;180:519–30.

[31] Grigorieff N. Frealign: An Exploratory Tool for Single-Particle Cryo-EM. Methods Enzymol. 2016;579:191–226.

[32] Mastronarde DN, Held SR. Automated tilt series alignment and tomographic reconstruction in IMOD. J. Struct. Biol. 2017;197:102–13.

[33] Jones TA, Zou JY, Cowan SW, Kjeldgaard M. Improved methods for building protein models in electron density maps and the location of errors in these models. Acta Crystallogr A. 1991;47:110–9.

[34] Afonine PV, Poon BK, Read RJ, Sobolev OV, Terwilliger TC, Urzhumtsev A, et al. Real- space refinement in PHENIX for cryo-EM and crystallography. Acta Crystallogr D Struct Biol. 2018;74:531–44.

[35] Chen VB, Arendall WB, 3rd, Headd JJ, Keedy DA, Immormino RM, Kapral GJ, et al. MolProbity: all-atom structure validation for macromolecular crystallography. Acta Crystallogr. D Biol. Crystallogr. 2010;66:12–21.

[36] Hunter JD. Matplotlib: A 2D graphics environment. Comput. Sci. Eng. 2007;9:90–5.

[37] Afonine PV. phenix.mtriage: a tool for analysis and validation of cryo-EM 3D reconstructions. Computational Crystallography Newsletter. 2017;8:25.

